# Sex-Specific Metabolic and Central Effects of GLP-1–Estradiol Conjugate in Middle-Aged Rats on a Standard or Western Diet

**DOI:** 10.1101/2025.04.07.647672

**Authors:** Jennifer E Richard, Ahmad Mohammad, Kimberly A Go, Andrew J McGovern, Rebecca K Rechlin, Tallinn FL Splinter, Stephanie E Lieblich, Lara K Radovic, Lydia Feng, Samantha A Blankers, Bin Yang, Jonathan D Douros, Brian Finan, Liisa AM Galea

**Affiliations:** Centre for Addiction and Mental Health, Toronto, ON Canada; Department of Physiology, Institute of Neuroscience and Physiology, The Sahlgrenska Academy at the University of Gothenburg, Gothenburg, Sweden; Djavad Mowafaghian Centre for Brain Health, University of British Columbia, Vancouver, BC, Canada; Institute of Medical Sciences, University of Toronto.; Department of Psychiatry, University of British Columbia, Vancouver, BC, Canada; Novo Nordisk Research Center Indianapolis, Inc. Indianapolis, IN USA; Department of Psychiatry, University of Toronto, Toronto, Canada

**Author notes:** Address all correspondence to: Liisa Galea, PhD, Treliving Family Chair in Women’s Mental Health, Senior Scientist, Centre for Addiction and Mental Health, Department of Psychiatry, University of Toronto.

**Keywords:** neurogenesis, hippocampus, insulin regulation, metabolic challenge, obesity, PSD-95, amygdala, associative learning, metabolic hormones, cytokines

## Abstract

Middle age represents a critical window for metabolic and cognitive health, particularly in the context of rising obesity and diabetes rates. Glucagon-like peptide-1 (GLP-1)-based therapies, which regulate blood glucose and body weight, show sex-specific effects, with estradiol potentiating their metabolic benefits. However, research on GLP-1’s cognitive and neuroprotective roles has largely been conducted in males. Here, we investigated the effects of GLP-1 conjugated to estradiol (GE2) on metabolism, cognition, cytokine levels and neurogenesis in the dentate gyrus of middle-aged male and female rats fed a standard (SD) or Western (WD) diet. In both sexes, WD increased body weight and plasma leptin levels, regardless of sex. GE2 treatment led to weight loss, enhanced cued and contextual fear memory, reduced cytokine levels in the hippocampus in SD rats, and increased neurogenesis in the dorsal dentate gyrus (DG), regardless of sex. Sex-specific differences were observed in fat distribution, glucose regulation, central cytokine levels, and neuroplasticity after WD and GE2 treatment. In females only, GE2 reduced visceral (gonadal) fat, reduced cytokines in the dorsal hippocampus, and improved basal blood glucose in response to a WD. In males only, GE2 restored neurogenesis in the DG after WD exposure, and reduced cytokine levels in the amygdala. These findings suggest that although WD increased body weight and GE2 improved associative learning in both sexes, both WD and GE2 had differential affects on metabolic hormones, insulin regulation, cytokine levels and neuroplasticity. Our findings underscore the importance of sex-specific approaches in metabolic and neuroprotective therapeutics in middle age.

## 1. Introduction

Diets high in saturated fats and refined carbohydrates are linked to obesity and diabetes (1) which increase the risk of dementia (2–4), likely through increased inflammation and insulin resistance (2–7), particularly in middle age (11–13). Indeed, diet-associated factors are included as modifiable risk factors for dementia, including diabetes, obesity, hypertension and high cholesterol (14). As the prevalence of obesity and diabetes has reached epidemic proportions in many regions throughout the world (15), such findings underscore the urgent need for targeted therapies addressing brain health in the face of high fat diets, especially during middle age - a period critical for metabolic and cognitive health.

Sex differences in obesity and diabetes are well established and influenced by both biological and hormonal factors. According to the 2025 Lancet Obesity report, the global prevalence of obesity is higher in females than in males, peaking around middle age in both sexes (16). Females typically accumulate more subcutaneous fat, whereas males accumulate more visceral fat—a pattern largely regulated by estrogens and their receptors (17). Following the menopausal transition, declining levels of ovarian hormones in females lead to a shift toward greater visceral fat accumulation, resembling the male pattern. Furthermore, the risk of developing diabetes is also affected by sex and menopause status, although diabetes is more prevalent in males overall, more females are living with the disease globally due to a reversal in prevalence after menopause (14). These differences highlight that middle age is a critical window of vulnerability for obesity and diabetes, both of which are risk factors for dementia.

With the growing rates of obesity and diabetes, glucagon-like peptide-1 (GLP-1)-based therapies have emerged as powerful tools to tackle these conditions (18). GLP-1 is an incretin hormone produced in the intestinal L-cells that promotes satiety, regulates blood glucose, activates brown fat metabolism and increases energy expenditure (19,20). Centrally, GLP-1 is primarily produced by neurons in the nucleus of the solitary tract (NTS) and modulates appetite and body weight through widespread neural projections (21,22). Intriguingly, the effects of GLP-1 on appetite reduction, glycemic control and body-weight reduction are more pronounced in females, at least in part due to estrogens (23–28). Sex differences are also observed in the neurological actions of GLP-1, where targeting certain brain areas produces more potent effects in females (e.g., the ventral tegmental area), whereas the opposite is true for other regions (e.g., the supramammillary nucleus and lateral hypothalamus) (29–31). However, the net outcome of central stimulation leads to more prominent reductions in appetite, blood glucose and body weight in adult females, suggesting that GLP-1 receptor treatments are more effective treatments for diabetes and obesity in females compared to males.

Beyond its role in metabolic regulation, GLP-1 can impact cognitive and emotional processes through GLP-1 receptors in the hippocampus, prefrontal cortex (PFC), and amygdala (32,33). Preclinical studies, primarily in males, demonstrate that GLP-1 receptor agonists improve hippocampus-dependent learning and memory under standard- and high-fat diet conditions, as well as in diabetes and AD models (34–39). Furthermore, GLP-1 receptor agonists have neuroprotective effects in the hippocampus, including enhanced neurogenesis in the dentate gyrus and reduced apoptosis in studies primarily conducted in adult males (34,40–43). The integrity of the hippocampus is important for spatial and episodic memory, and it is one of the first brain regions affected in AD (44). GLP-1 receptor agonists are associated with lower dementia rates and improved cognitive outcomes in clinical studies, independent of metabolic improvements (39,45). Thus, evidence suggests that GLP-1 receptor agonists have a neuroprotective role and can enhance learning and memory, although there is limited research examining effects in females or in middle-aged adults.

Middle age is a critical period for diet-induced cognitive deficits (11–13), coinciding with the menopausal transition in females. This transition is characterized by declining ovarian hormone levels, including estrogens, which play a neuroprotective role during certain circumstances (46). Estradiol (E2)-based hormone therapies can improve cognition in postmenopausal women (47–51). E2 also promotes hippocampal plasticity (52–55) through several possible actions (56), including through its actions on synaptic proteins such as PSD-95 (57,58), and reduced proinflammatory cytokines in the hippocampus (59,60) in female rodents. Moreover, E2 potentiates GLP-1’s effects on food intake in female rats (61) and reward (62) in both sexes, but to a greater extent in females compared to males, suggesting a synergistic relationship between these hormones at least in young adult rats. But to date, no studies have examined the effects of combined GLP and E2 treatment on hippocampus-dependent memory and cytokine levels in the hippocampus in both sexes at middle age.

To harness the cognitive and neuroprotective benefits of both hormones, we utilized GE2 (63), a conjugate combining GLP-1 and E2. This study investigated whether GE2 improves cognition, reduces central cytokine levels, and enhances synaptic plasticity in the hippocampus of middle-aged male and female rats fed a standard (SD) or high-fat, high-sugar (Western) diet (WD). We hypothesize that GE2 treatment will mitigate the cognitive and inflammatory impacts of a WD and enhance cognition, reduce cytokine levels in brain regions involved in associative memory, and increase neuroplasticity in limbic regions of the brain in middle-aged male and female rats. Furthermore, whereas sex-specific effects of GLP-1 have been reported for its metabolic outcomes and reward behaviors (30,31,62), research on GLP-1’s effects on cognition and neuroprotection have been conducted mainly in males. This research addresses a significant gap by examining these outcomes in both male and female rats. We hypothesize that more pronounced effects will be observed in females due to the potentiating effects of estradiol conjugated to GLP-1.

## 2. Methods

### 2.1. Animals

Adult male (n=57) and female (n=56) Sprague–Dawley rats were bred in house from breeding pairs obtained from Charles River, Quebec, Canada. Twenty-nine males and 30 females were used for the main experiment and received either GE2 or control, whereasthe remaining ratsreceived equimolar doses of GLP-1 or E2. The data for the latter groups are included in the Supplement. After weaning rats were double-housed in a 12-h light/dark cycle. Rats had *ad libitum* access to water and SD (Picolab®, rodent diet 5053) until 10 months of age in which they were divided into weight-matched groups within each sex, and fed a SD (Picolab®, rodent diet 5053) or WD (Research Diets (RD) Western Diet, D12079B, Research Diets Inc., New Brunswick, NJ, USA) for the following 8 weeks of the experiment as described below and demonstrated in Figure 1A. All protocols were approved by the Institutional Animal Care Committee at the University of British Columbia and conformed to the guidelines set out by the Canadian Council on Animal Care.

**Figure 1.**
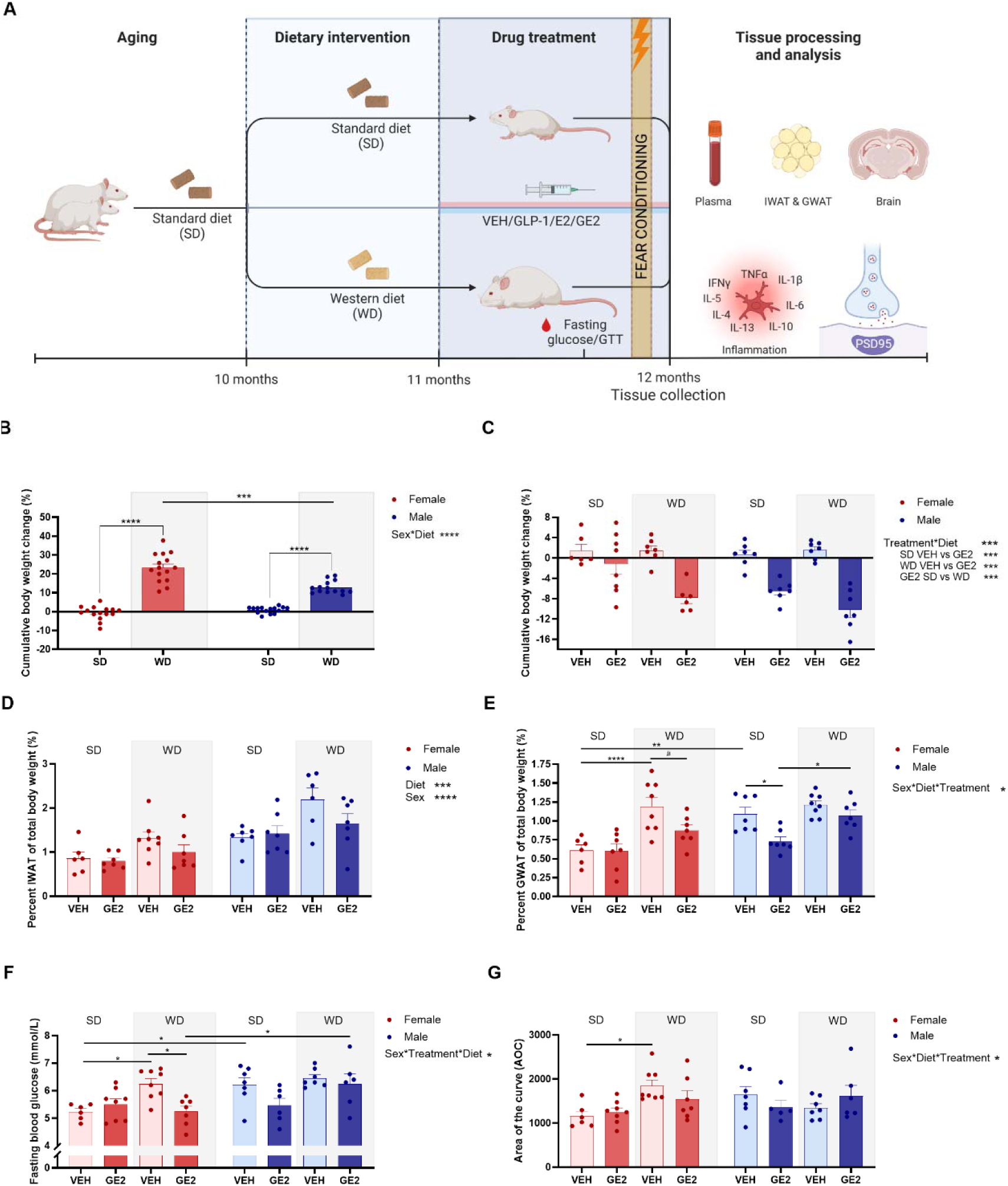
Diet and treatment affect body weight, fat and blood glucose in a sex-specific manner. Graphical illustration of experimental plan (A). Mean (±SEM) percent body weight (B), as well as mean (±SEM) percentage of inguinal white adipose tissue (IWAT; D) and gonadal white adipose tissue (GWAT; E) after 4 weeks of treatment. Mean (±SEM) Basal blood glucose levels after an overnight fast (F) and area of the curve (AOC) after being challenged by a glucose tolerance test (GTT; G). * p < 0.05, ** p < 0.01, *** p < 0.001, **** p < 0.0001 and # p < 0.100. Diet only: n = 33-34 (SD) and n = 34-35 (WD); Treatment groups: n = 6-8 (VEH) and n = 5-8 (GE2) per diet and sex. VEH = vehicle; GE2 = GLP-1-E2; SD = standard diet; WD = western diet. Timeline image created using BioRender.com.

### 2.2. Drugs

Compound GE2 stable conjugate and GLP-1 (GLP-1-X) were provided by Brian Finan and Bin Yang (Novo Nordisk Research Center Indianapolis, Indianapolis, IN, USA). Detailed information on the pharmacokinetic profile for the stable conjugate and GLP-1 have previously been reported in (63). A dose of 20 µg/kg (4.5nmol/kg) was chosen based on (63,64) and under guidance from the Novo Nordisk team. This dose is 2-6 times lower than previous studies using the conjugate (63,65) to minimize the risk of side effects; incidences of which have been higher in females (29). GE2 was dissolved in saline (0.9% sodium chloride saline solution for injection, JB1323, Baxter); used as vehicle.

### 2.3. Experimental timeline

At 11 months of age (after one month of diet exposure) females and males received daily subcutaneous injections of vehicle (saline) or GE2 as described above (n = 6-7 (vehicle) and n = 6-8 (GE2); Fig. 1A). Rats were weighed weekly during the diet only period and daily prior to drug injection. Starting at 11 months, or 4 weeks after diet manipulation, male and female rats received daily subcutaneous injections of GE2, or vehicle (saline) for 1 month until tissue collection. Vaginal samples were collected via lavage during the diet and treatment periods. Eighty percent of females were irregular or acyclic during diet and treatment manipulations (**Supplementary Fig. 14**). In separate studies, we also examined the individual contribution of GLP-1 and E2 alone to determine the relative contribution of these hormones to compare to the conjugate, and data for all treatment groups are available in the supplementary section.

### 2.4. Glucose tolerance test (GTT)

GTT was performed after 20 days of hormone/vehicle treatment. Rats were fasted overnight (from 8 pm to 9 am), then weighed and the tip of the tail was scored using a fresh scalpel. Blood was collected from the tail vein and basal blood glucose was measured using a glucometer (OneTouch Ultra®2, Lifescan). Rats then received an intraperitoneal injection of glucose solution (2g/kg body weight with a volume of 1mL/kg). Blood was sampled by gently massaging the tail, 15, 30, 60, and 120 minutes after injection and blood glucose measured using the glucometer. Area of the curve (AOC) was calculated for each subject by subtracting the starting glucose value from the value at each time point as described in (66).

### 2.5. Contextual and cued fear conditioning

After 1 month of treatment (2 months of diet), rats were tested using the fear conditioning paradigm, which relies on the integrity of the amygdala (cued fear conditioning), and hippocampus (contextual fear conditioning) (67). The apparatus and testing paradigm was previously described in (68). Briefly, conditioning and testing were conducted in four identical observation chambers (30.5 × 24 × 21 cm; Med Associates, St Albans, VT), illuminated by a 100 mA house light and enclosed within sound-attenuating boxes. A speaker connected to a programmable audio generator (ANL-926, Med Associates) provided tones (4 kHz, 80 dB) and 19 stainless steel rods, spaced 1.5 cm apart, were connected to a shock generator and scrambler for the delivery of an unconditioned stimulus foot shock (1.0 mA). A video camera positioned above each chamber recorded behavior for video scoring. Ventilation fans in each box supplied background noise (70 dB, A scale). Locomotor activity was measured (Med Associates) and time spent freezing was scored manually using video footage with the experimenter blinded to group membership.

Training and testing were conducted over 2 days. On day 1 (conditioning day) rats were injected with GE2 or vehicle by one experimenter. Thirty minutes later, rats were individually transported to the testing room in a clean, unused, transparent cage, on a four-wheeled cart by a different experimenter. Rats were then placed in the Context A operant chamber. Context A consisted of bare aluminum and Plexiglass walls, and a lavender air freshener was used as an odor cue. Three minutes after placement into the chamber, rats were given three presentations of the tone conditioned stimulus (CS; 30 seconds), each co-terminating with a foot shock (2 seconds) with 60 second intershock intervals. Sixty seconds after the third and final tone/shock pairing (after 3 minutes of exposure to Context A), rats were immediately returned to their home cage. To assess whether there were group differences in pain sensitivity, we recorded freezing behavior during the minute after each shock presentation.

On day 2 (24 h after conditioning), rats were again given hormone treatment and then individually transported, in a clean, unused, transparent cage, back to the testing room 30 minutes after injection. The same procedure was followed as the conditioning day to activate the representation of the context. Rats were placed back into Context A (same as described above) for 8 min to assess contextual fear conditioning (no tones or shocks were present). One hour later, rats were transported back to the testing room in an unused, transparent cage, but along a completely novel route. Rats were placed into the novel Context B, which had striped inserts covering the Plexiglass walls, and a tropical air freshener. Three minutes after being placed in the box, they were presented with three tones (4kHz, 80 dB, 30 seconds), 60 seconds apart. Time spent freezing to the tone (cue) was measured during each tone presentation to assess cued fear conditioning. Unfortunately, due to technical difficulties in two of the boxes, the tone did not play during conditioning for 14 animals, and they were therefore excluded from the analysis.

### 2.6. Tissue and blood collection

Trunk blood was collected following live decapitation and placed into cold EDTA-coated tubes. Tubes were centrifuged for 15 minutes, and plasma was stored at −80°C. Brains were quickly extracted, and the hemispheres were divided. The left hemisphere was flash frozen on dry ice and stored at −80°C. Brain areas of interest (amygdala, dorsal and ventral hippocampus, PFC) were punched from slices (50µm sections) using a cryostat (−20°C), and cytokine levels analyzed using electrochemiluminescence immunoassay kits. The right hemisphere was placed in paraformaldehyde (PFA) and stored at 4°C. After 24 hours the brains were transferred to a 30% sucrose/PBS solution for cryoprotection. Serial coronal sections (30 µm) were cut with a freezing microtome (SM2000R; Leica, Richmond Hill, ON) across the extent of the hippocampus (collected in 10 series) and stored in an antifreeze solution (containing 0.1M PBS, 30% ethylene glycol, and 20% glycerol) and stored at −20°C for immunohistochemistry. Inguinal (IWAT) and gonadal (GWAT) white adipose tissue were collected and weighed, and a sample was collected and flash frozen on dry ice.

### 2.7. Hippocampal cytokine measurements

Electrochemiluminescence immunoassay kits (Meso-Scale Discovery; Rockville, MD) were used to measure cytokines (IFN-γ, IL-1β, IL-4, IL-5, IL-6, IL-10, IL-13, CXCL-1, and TNF-α; V-PLEX Proinflammatory Panel 2 Rat Kit; Cat# K15059D) and postsynaptic density 95 (PSD95; PSD95 kit; Cat# K150QND) in homogenized amygdala, hippocampus (dorsal and ventral) and PFC samples. PSD95 is a postsynaptic protein that anchors glutamatergic synapases (69) and was chosen because estradiol influences PSD-95 in the hippocampus (58) and is related to associative learning (70,71). Each sample was homogenized using a TissueLyser III (Qiagen) with 5 homogenization beads in 200 µl of cold Tris lysis buffer (150 mM NaCl, 20 mM Tris, pH 7.4, 1 mM EDTA, 1 mM EGTA, 1% Triton X) containing a cocktail of protease (cOmplete Protease Inhibitor Cocktail, Roche) and phosphatase inhibitors (Phosphatase Inhibitor Cocktail 2 and 3, Sigma Aldrich), NaF and PMSF in DMSO. Homogenates were centrifuged at 100 rpm for 30 seconds at 4°C and supernatants were aliquoted and stored at −80°C. Protein concentrations were determined using a BCA protein assay kit (Thermo Scientific). Homogenates were diluted with Tris lysis buffer 1:10 for PSD95 and 1:4 for proinflammatory analysis. Samples were run in duplicate following the manufacturer’s protocols, with the exception of longer sample and calibrator incubation for cytokine plates (overnight instead of 2 hours at 4°C). Plates were read with a MESO QuickPlex SQ 120 (Meso Scale Discovery) and data were analyzed using the Discovery Workbench 4.0 software (Meso Scale Discovery). Cytokine levels were obtained using a standard curve and expressed as pg/mg protein and PSD95 measured protein signal levels which were normalized to protein and expressed as signal/ pg mg^-1^.

### 2.8. Plasma hormone levels

U-PLEX Metabolic Group 1 Rat Kits (Meso-Scale Discovery; Rockville, MD; Cat# K15200L) were used to measure metabolic hormones (c-peptide, ghrelin, glp-1, glucagon, insulin, leptin and PYY) in plasma. Plasma was diluted using a 2-fold dilution using the Metabolic Assay Working Solution. Samples were run in duplicate following the manufacturer’s protocol. Plates were read with a MESO QuickPlex SQ 120 (Meso Scale Discovery) and data were analyzed using the Discovery Workbench 4.0 software (Meso Scale Discovery). Hormone levels were obtained using a standard curve and expressed as pg/ml.

### 2.9. Immunohistochemistry

One series of hippocampal sections from each rat, collected using the microtome as described in section 2.6, was added to 12-well tissue culture plates with 0.1 M phosphate-buffered saline (PBS) and rinsed 3 times at room temperature using nets for 10 minutes each. The tissue was then incubated in 10% Triton X for 30 minutes. Sections were added to the blocking solution (10% normal donkey serum (NDS) and 3% Triton X in PBS) for 1 hour. The tissue was immediately incubated in rabbit anti-doublecortin (DCX; 4604S, Cell Signal Technology Inc. Danvers) diluted 1:1000 in blocking solution (10% NDS and 3% Triton X in PBS) for ∼48 hours at 4°C. Following PBS rinses (10 minutes X3), tissue was incubated for 4 hours in Alexa Fluor 488 conjugated donkey anti-rabbit (1:500; A-21206, Thermo Fisher Scientific) in a solution PBS containing 10% NDS, 3% Triton-X. Sections were then washed in PBS for 3x 10 minutes and then stained with 4,6-diamidino-2-phenylindole (DAPI; 1:500; D1306, Thermo Fisher Scientific) for 2.5 minutes at room temperature. The tissue was rinsed for 5 minutes (3X). Finally, tissue was mounted onto Superfrost/Plus slides (Fisher scientific) and cover slipped using PVA-DABCO mounting medium.

### 2.10. DCX-expressing cells: quantification and microscopy

DCX is a microtubule-associated protein expressed in immature neurons, and it is commonly measured in the dentate gyrus (DG) as a marker of adult neurogenesis (72). DCX-expressing cells were imaged with Zeiss Axio Scan.Z1 (Carl Zeiss Microscopy, Thornwood, NY, USA) with a 20x objective using fluorescent imaging. The area around the DG was first identified and measured using DAPI only and all DCX-expressing cells within the marked area were then quantified by an investigator blinded to experimental conditions. Two dorsal and two ventral sections were quantified for each rat. The density was calculated by dividing the amount of DCX-expressing cells with the area of the DG for each section. Photomicrographs were acquired using the Fluoview 4000 confocal microscope (Evident Scientific) with a 20x objective lens. DAPI was visualized using the 405nm laser (blue) and DCX was visualized using the 488nm laser (green). Contrast was adjusted to remove background signal and single cell images were deconvolved using cellSens imaging software.

### 2.11. Statistical analyses

All the data are presented as mean ± standard error of the mean. Data were analyzed using TIBCO Statistica software (v. 9, StatSoft, Inc., Tulsa, OK, USA) using analyses of variance (ANOVA) with sex (male, female), diet (SD, WD) and treatment (vehicle, GE2) as between subject factors on dependent variables of interest, and plotted using GraphPad Prism (GraphPad Software, Inc., San Diego, CA). Effect sizes (partial eta squared (_p_^2^) and Cohen’s *d*) are reported for significant effects. Post hoc comparisons used Newman-Keul’s and *a priori* comparisons were subjected to a Bonferroni correction. P-values ≤ 0.050 were considered statistically significant, and p-values between 0.05-0.100 are reported as trends.

#### 2.11.1. For z-stack and PERMANOVA

*Data Preprocessing and Imputation:* To address missing values in our dataset, we employed a multiple imputation approach using the random forest (RF) method implemented in the mice package (73). A predictor matrix was constructed establishing region-specific relationships, giving full weight to predictors within the same brain region (dHPC, vHPC, Amyg, PFC), partial weight (0.5) to predictors from other regions, and including demographic variables (Group, Treatment, Diet) as full predictors. The RF imputation was executed with 5 multiple datasets (m = 5), 20 iterations, and 200 trees per forest. The quality of imputation was assessed through density plots comparing original and imputed distributions for each variable within each brain region, ensuring the preservation of the original data structure (**Supplementary Fig. 5**).

#### 2.11.2. Regional Cytokine Profiling and Multivariate Analysis

The multivariate patterns of cytokine expression across brain regions were analyzed using a permutational multivariate analysis of variance (PERMANOVA) approach implemented in the R vegan package on imputed data (74,75). For each region, we extracted cytokine measurements and constructed Euclidean distance matrices after standardizing the data (z-scoring). Cytokines were categorized as either TH1 (pro-inflammatory: IL1β, TNFα, IFNγ) or TH2 (anti-inflammatory: IL4, IL5, IL10, IL13) to characterize distinct immune response patterns. The PERMANOVA was performed using 15,000 permutations and a three-way factorial design (Sex × Diet × Treatment). Non-metric multidimensional scaling (NMDS) was used to visualize multivariate patterns in cytokine profiles. Mean inflammatory profiles were calculated by averaging z-scored values within TH1 and TH2 categories for each region. Differences between inflammatory profiles within experimental groups were assessed using pairwise Wilcoxon tests with false discovery rate (FDR) correction.

Software and Packages: All analyses were conducted in R (version 4.3.2). The following packages were utilized: mice for multiple imputation; tidyverse for data manipulation and visualization and vegan for multivariate analyses.

## 3. Results

### 3.1. WD increased, whereas GE2 reduced, body weight in both sexes

As expected, four weeks of WD consumption resulted in weight gain, compared to SD in both sexes (males SD vs WD: *p* < 0.001, Cohen’s *d* = 4.576; and females SD vs WD: *p* < 0.001, Cohen’s *d* = 4.049; **Fig. 1B**), although females gained significantly more weight than males (Sex and Diet interaction: F(_1, 55_) = 25.237, □_p_^2^ = 0.315, *p* < 0.001).

After one month of the hormone treatment, GE2-treated subjects of both sexes lost significantly more weight their vehicle-treated counterparts (Treatment and diet interaction: F(_1,47_) = 9.936, □_p_^2^ = 0.175, *p* = 0.003; SD: *p* = 0.001, Cohen’s *d* = 1.140; WD: *p* = 0.000, Cohen’s *d* = 3.733). Not surprisingly, GE2 treatment led to a larger body-weight reduction on WD than SD fed (*p* = 0.000, Cohen’s *d* = 1.238). The main effects of Treatment (F(_1, 47_) = 72.550, □_p_^2^ = 0.607, *p* = 0.000), Diet (F(_1, 47_) = 6.798, □_p_^2^ = 0.126, *p* = 0.012) and Sex (F = 5.451, □_p_^2^ = 0.104, *p* = 0.024), were also significant, however, there were no other significant main or interaction effects (*p* ≥ 0.062).

### 3.2. Diet and GE2 treatment affect GWAT in a sex-specific manner

GWAT was collected as a measure of visceral fat, a type of fat that is more detrimental to health, increasing the risk of metabolic disorders and cardiovascular disease, whereas IWAT was collected as a measure of subcutaneous fat, which is considered a metabolically healthy fat type (76). WD increased IWAT compared to SD (main effect of diet: F(_1, 46_) = 13.735, □_p_^2^ = 0.230, *p* = 0.001, **Fig. 1D**), especially in males. Males also had more IWAT than females overall, regardless of diet (main effect of sex: F(_1, 46_) = 30.170, □_p_^2^ = 0.396, *p* < 0.001).

WD consumption led to a significant increase in GWAT (measure of visceral fat) compared to SD, in vehicle-treated groups, but only in females (*p* < 0.000, Cohen’s *d* = 2.147; Sex x Diet x Treatment interaction: F(_1, 47_) = 4.485, □_p_^2^ = 0.087, *p* = 0.040, **Fig. 1E**). There was also a strong trend towards a reduction in GWAT in WD-fed females treated with GE2 compared to vehicle-treated females (*p* = 0.057). Furthermore, GE2 significantly reduced GWAT in SD-fed males compared to vehicle-treated SD males (*p* = 0.028, Cohen’s *d* = 1.848). GE2-treated males had more GWAT on the WD than the SD (*p* = 0.024, Cohen’s *d* = 0.099). In addition, vehicle-treated males had significantly more GWAT than vehicle-treated females on the SD (*p* = 0.003, Cohen’s *d* = 2.324). There were no other significant main effects or interactions (all *p*’s ≥ 0.071).

### 3.3. Diet and GE2 treatment affect fasting blood glucose in a sex-specific manner

Next, we investigated the effects of diet and hormone treatment on blood glucose. WD consumption increased fasting blood glucose in vehicle-treated rats compared to SD, however, only in females (*p* = 0.035, Cohen’s *d* = 2.342; Sex x Diet x Treatment interaction: F(_1, 47_) = 6.624, □_p_^2^ = 0.124, *p* = 0.013, **Fig. 1F**). GE2 treatment in WD females significantly reduced fasting blood glucose compared to vehicle (*p* = 0.032, Cohen’s *d* = 2.002); an effect not present in males (*p* = 0.803). Furthermore, GE2-treated WD females had lower fasting blood glucose levels than GE2-treated WD males (*p* = 0.024, Cohen’s *d* = 7.084). Furthermore, vehicle-treated males on SD had higher fasting glucose than vehicle-treated females on SD (*p* = 0.022, Cohen’s *d* = 1.869). The main effects of Treatment (F(_1, 47_) = 5.720, □_p_^2^ = 0.109, *p* = 0.021), Diet (F = 5.513, □_p_^2^ = 0.105, *p* = 0.023) and sex (F = 13.599, □_p_^2^ = 0.224, *p* = 0.001), were also significant, there were no other significant main or interaction effects (*p >* 0.139).

After a glucose challenge, we found an increase in the area under the curve (AUC) in WD-fed vehicle-treated females compared to SD-fed vehicle-treated females (*p* = 0.050; Sex x Diet x Treatment interaction: F(_1, 46_) = 4.663, □_p_^2^ = 0.092, *p* = 0.036, **Fig. 1G**), but not in males (*p* = 0.650). There was also a significant interaction between sex and diet (F(_1, 46_) = 5.516, □_p_^2^ = 0.107, *p* = 0.023) and a significant main effect of diet (F(_1, 46_) = 4.618, □_p_^2^ = 0.091, *p* = 0.037), but no other significant main or interaction effects (*p >* 0.627).

### 3.4. GE2 treatment increases plasma GLP-1 only in females and PYY in both sexes, WD increased leptin levels

To investigate if the rodents were metabolically affected by WD consumption after 8 weeks, we measured the levels of several plasma metabolic hormones. First, we wanted to know if our treatment affected the levels of plasma GLP-1. We found that GE2 treatment significantly increased plasma GLP-1 compared to vehicle in females (*p* = 0.001; Cohen’s *d* = 0.157; sex and treatment interaction: F(_1, 46_) = 7.146, □_p_^2^ = 0.134, *p* < 0.010; **Fig. 2A**), but not in males (*p* = 0.806). The main effect of treatment (F(_1, 46_) = 9.137, □_p_^2^ = 0.166, *p* = 0.004) and sex (F = 4.586, □_p_^2^ = 0.091, *p* = 0.038) were also significant, but there were no other significant main effects or interactions (all *p* ≥ 0.105).

**Figure 2.**
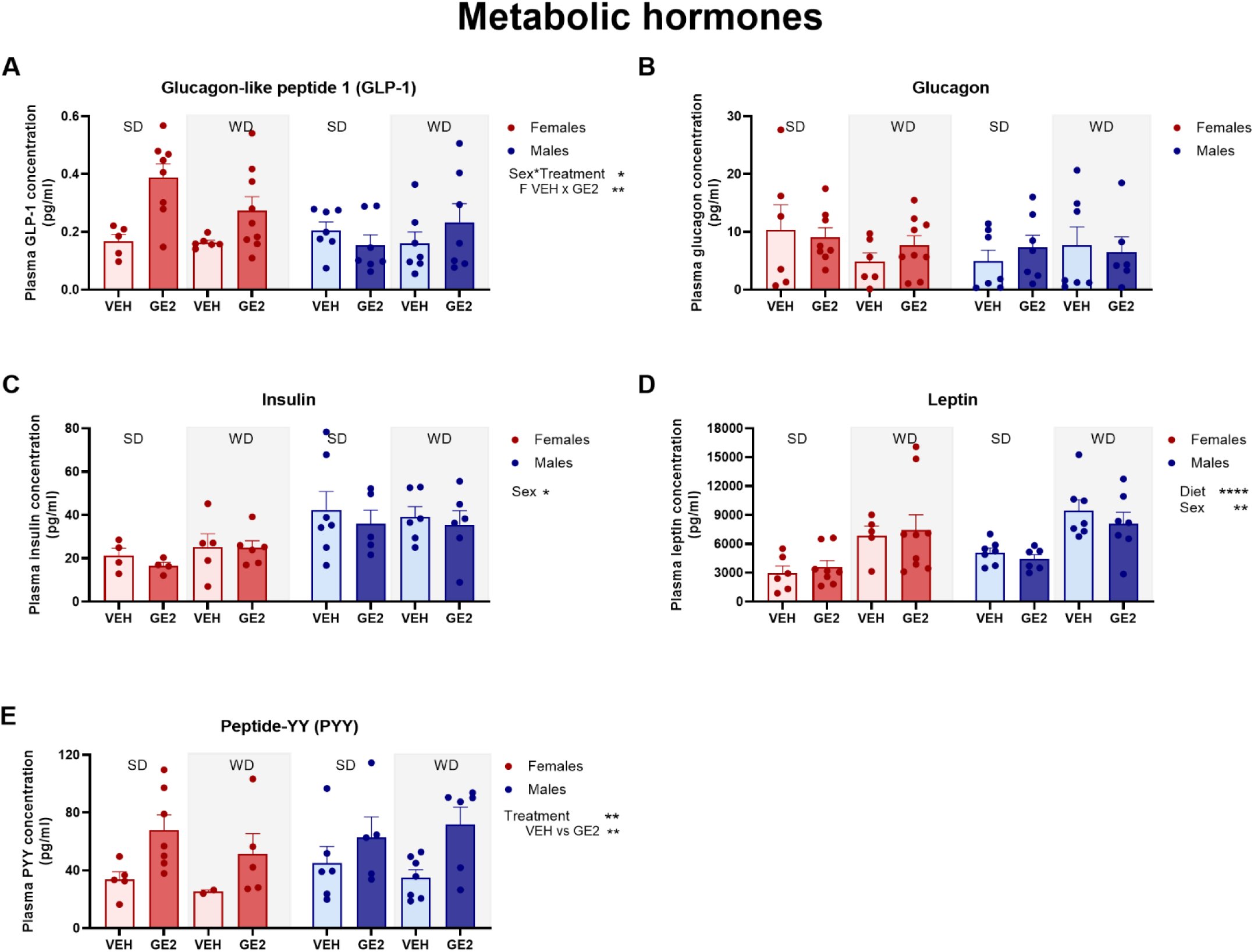
GE2 treatment increases plasma peptide YY (PYY) levels in both sexes, and GLP-1 levels only in females, compared to Vehicle. Mean (±SEM) levels of glucagon-like peptide-1 (GLP-1; A), glucagon (B), insulin (C), leptin (D) and PYY (E) in plasma. Main and interaction effects are listed with graphs* p < 0.05 ** p < 0.01, **** p < 0.0001. n = 2-7 (Vehicle) and n = 5-9 (GE2) per diet and sex. SEM = standard error of the mean.

Next, we looked at glucagon and insulin, hormones regulated by GLP-1. We saw no statistical differences in glucagon levels by sex, treatment or diet, nor an interaction between these factors (all *p* ≥ 0.240, **Fig. 2B**). Males demonstrated higher levels of insulin than females (main effect of sex: F(_1, 37_) = 5.576, □_p_^2^ = 0.131, *p* = 0.024; **Fig. 2C**), but there were no other significant main effects or interactions (all *p*’s ≥ 0.155).

Leptin is directly affected by fat content. As expected, leptin levels were increased in WD compared to SD fed subjects (main effect of diet: (F(_1, 46_) = 21.606, □_p_^2^ = 0.320, *p* < 0.001; **Fig. 2D**). Males had higher levels of leptin than females (main effect of sex: F(_1,46_) = 7.280, □_p_^2^ = 0.137, *p* = 0.010). There were no other significant main effects or interactions (all *p*’s ≥ 0.399).

Levels of PYY, an anorexigenic peptide, were increased in GE2-treated subjects of both sexes (main effect of treatment: F(_1,34_) = 10.896, □_p_^2^ = 0.243, *p* = 0.002; vehicle and GE2: *p* = 0.002; Cohen’s *d* = 1.131; **Fig. 2E**). Plasma ghrelin and c-peptide were also measured but most values were below detection level (all ghrelin levels were below detection, as expected since the rodents did not undergo long-term fasting prior to blood collection to ensure that GLP-1 would be at detectable levels, and >80% of c-peptide levels were below detection), and thus we were unable to statistically analyze these hormones.

### 3.5. GE2 treatment improved both cued and contextual fear memory in both sexes

For cued fear conditioning, there was a strong trend for the WD to reduce freezing in the cued fear conditioning task (main effect of diet: F(_1,33_) = 4.037, □_p_^2^ = 0.109, *p* = 0.053), and a priori comparisons showed that WD fed males had reduced freezing compared to SD-fed males (p=0.019, Cohen’s d=1.13) which was not observed in females (p=0.911). GE2-treated rats displayed more time freezing than vehicle-treated rats in both sexes (males: *p* < 0.001, Cohen’s *d* = 1.899; females: *p* = 0.043, Cohen’s *d* = 1.240), despite the sex and treatment interaction (F(_1,33_) = 4.468, □_p_^2^ = 0.119, *p* = 0.042; **Fig. 3C**) in the cued fear conditioning task. The main effects of treatment (F(_1,33_) = 30.065, □_p_^2^ = 0.477, *p* < 0.001) and sex (F = 35.776, □_p_^2^ = 0.149, *p* = 0.022) were also significant. As for the contextual fear conditioning task, GE2-treated rats displayed more time freezing to the context than vehicle-treated rats regardless of sex (main effect of treatment: F(_1,48_) = 39.617, □_p_^2^ = 0.452, *p* < 0.001; **Fig. 3B**). No other significant main effects or interactions for contextual fear conditioning were present (all *p* ≥ 0.353).

**Figure 3.**
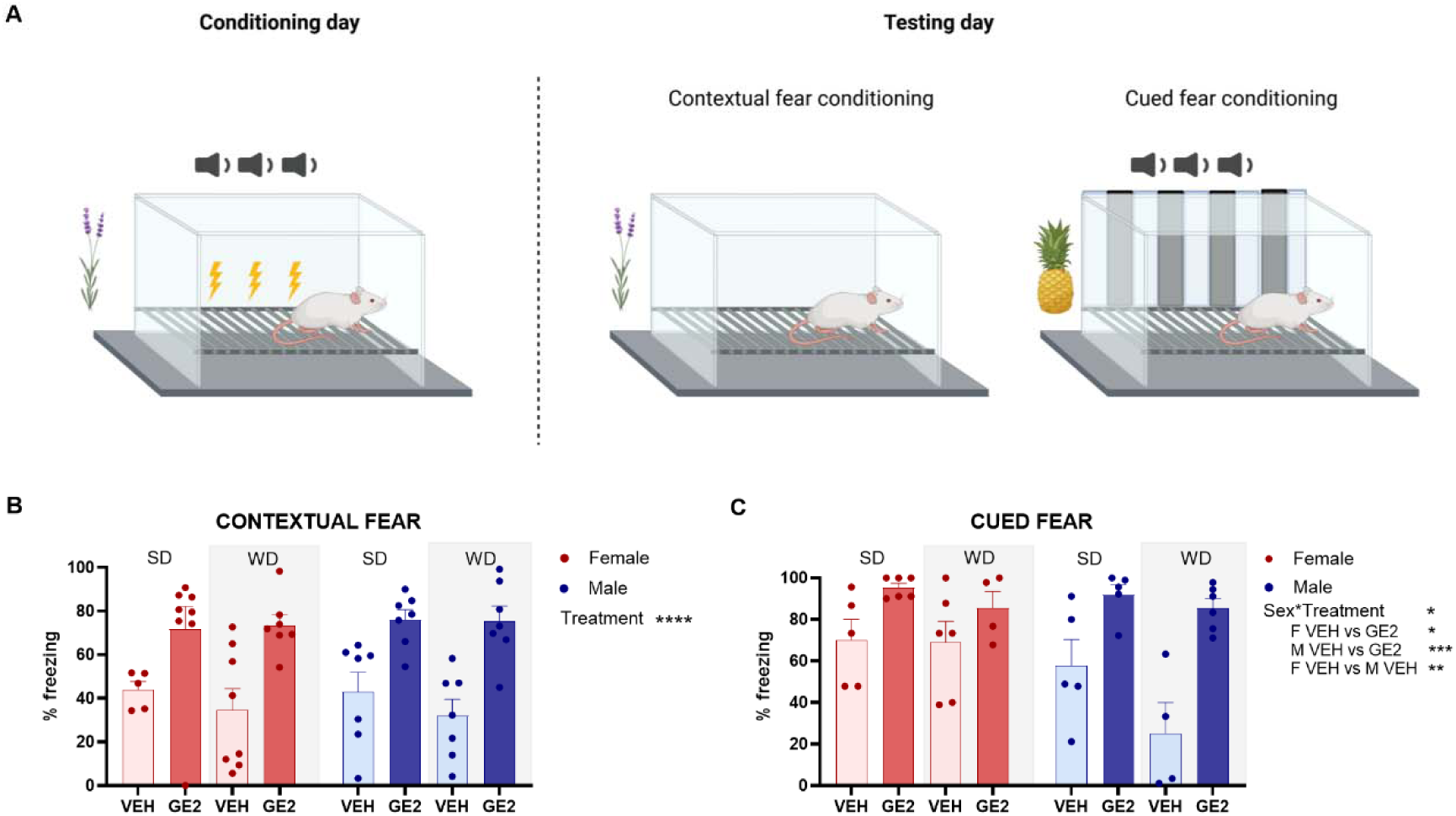
GE2 increases freezing in both the contextual and cued fear conditioning tests. Schematic representation of the 2-day fear conditioning training and testing (A). Mean percent freezing (±SEM) during the contextual fear conditioning task (B) and the cued fear conditioning task (C). * p < 0.05, ** p = 0.01, *** p < 0.001 and **** p < 0.0001. n = 5-10 (VEH) and 4-7 (GE2) per diet and sex. F = female; M = male, VEH = vehicle; GE2 = GLP-1-E2, SD = standard diet; WD = western diet; SEM = standard error of the mean. Schematic representation of fear conditioning task created using BioRender.com.

### 3.6. GE2 treatment increases neurogenesis in the dorsal DG depending on sex and diet

WD reduced the density of DCX-expressing cells compared to SD (*p* = 0.024, Cohen’s *d* = 1.309), but only in males, and GE2 treatment normalized the density of DCX-expressing cells under a WD in males (*p* = 0.008, Cohen’s *d* = 4.023; **Fig. 4C**). In females, GE2 treatment also increased the density of DCX-expressing cells compared to vehicle treatment, but only in the SD condition (a priori comparison: *p* = 0.036, Cohen’s *d* = 0.725; sex, diet and treatment interaction: F(_1,46_) = 3.445, □_p_^2^ = 0.070, *p* = 0.070). The main effect of treatment was also significant (main effect of treatment: F(_1,46_) = 8.883, □_p_^2^ = 0.162, *p* = 0.005). There were no other significant main effects of interactions in the dorsal DG (all *p* ≥ 0.092). In the ventral DG, we found no significant main effects or interactions (all *p* ≥ 0.334; **Fig. 4D**).

**Figure 4.**
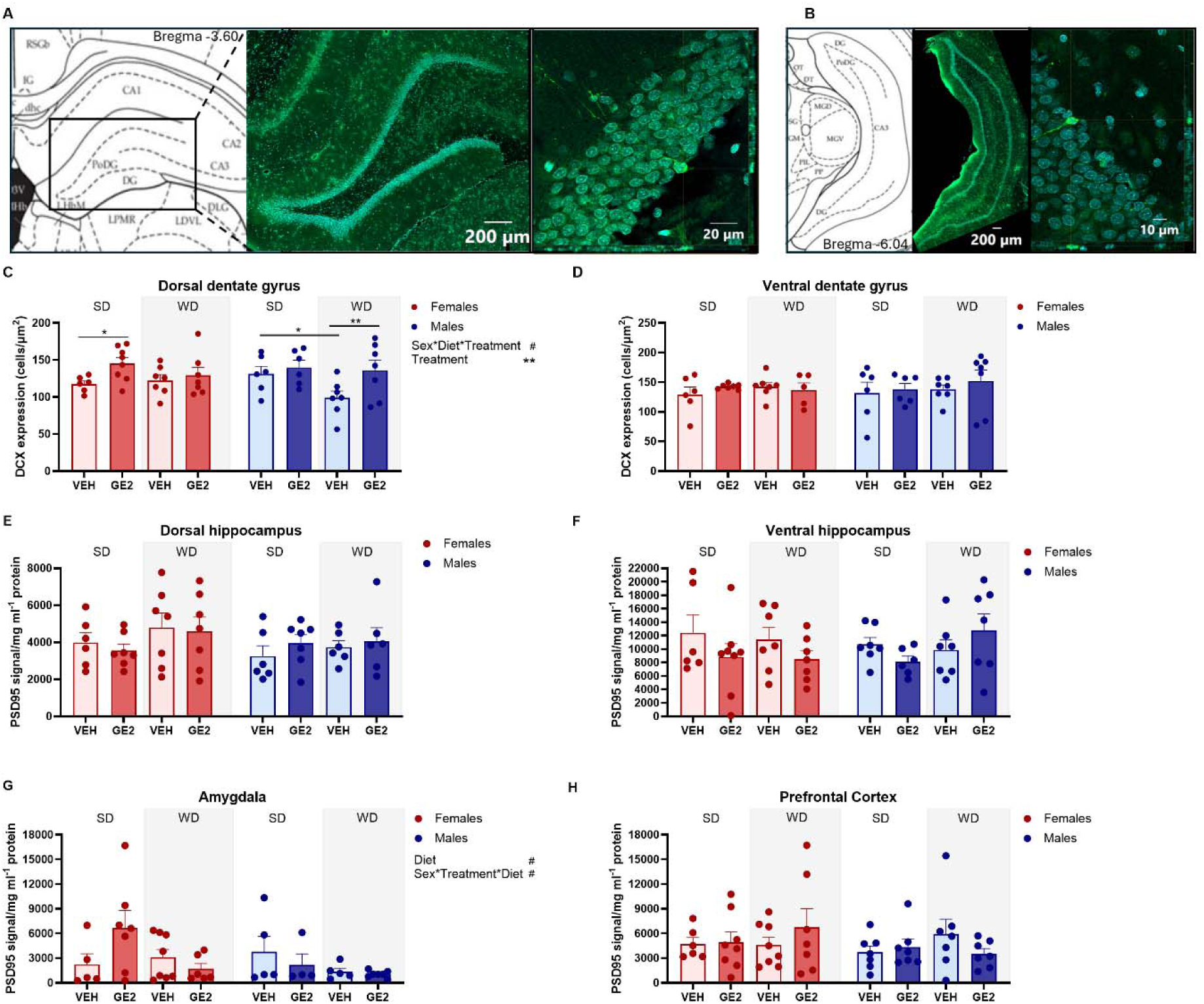
WD reduces doublecortin (DCX) expression in the dorsal dentate gyrus (DG) and GE2 treatment increases the density of DCX in the dorsal DG in SD females and WD males, whereas postsynaptic density 95 (PSD95) signal is unchanged. Representative image of DCX-expressing cells (green) in the dorsal (A) and ventral (B) DG. Mean DCX cells per µm^2^ (±SEM) measured in the dorsal (C) and ventral (D) DG. PSD95 signal in the dorsal hippocampus (E) ventral hippocampus (F), amygdala (G), and prefrontal cortex (H). DCX is depicted in green and DAPI in light blue. # p < 0.100, ** p < 0.010. n = 5-8 (vehicle) and 6-8 (GE2) per diet and sex. SD = standard diet; WD = western diet; SEM = standard error of the mean.

### 3.7. PSD95 levels were not significantly affected by sex, diet or GE2 treatment across brain regions

Levels of the postsynaptic protein PSD95 were not affected by diet, sex or treatment in the dorsal hippocampus (all *p* ≥ 0.161; **Fig. 4E**), ventral hippocampus (all *p* ≥ 0.182; **Fig. 4F**), or PFC (all *p* ≥ 0.202; **Fig. 4H**). In the amygdala, there was a trend towards lower levels of PSD95 in WD compared to SD subjects (main effect of diet: F(_1,38_) = 3.838, □_p_^2^ = 0.092, *p* = 0.057; **Fig. 4G**). Furthermore, there was a trend towards an interaction between sex, diet and treatment in the amygdala (main effect of diet: F_(1,38)_ = 3.263, □_p_^2^ = 0.079, *p* = 0.079).

### 3.8. Chronic WD consumption did not result in wide-scale increases in central cytokine levels, but did increase the levels of certain cytokines in a sex- and region-specific manner, and GE2 reduced inflammation in the hippocampus in SD-fed rats, as well as in the amygdala of males regardless of diet

Brain aging is associated with increased cytokine levels, however diets, such as the WD, may further affect cytokine levels, however, in the current study, we did not find wide-scale cytokine levels in the brain regions measure. We did, however, find increases in the levels of certain cytokines in response to WD in a sex-dependent manner, in which females demonstrated increased cytokine levels in the amygdala, whereas males trended towards increased levels in the ventral hippocampus. GE2 treatment reduced cytokine levels in the dorsal and ventral hippocampus, however only in SD subjects, regardless of sex, as well as in the amygdala in males regardless of diet.

#### 3.8.1. Dorsal hippocampus

GE2 treatment reduced the levels of certain cytokines in the dorsal hippocampus (**Fig. 5**). Z-score analysis of the cytokine profiles in the dorsal hippocampus revealed that GE2 reduced cytokine levels in SD rats (F_(1,25)_ = 8.971, R^2^ = 0.264, *p* < 0.001; diet and treatment interaction: F_(1,54)_ = 5.225, R^2^ = 0.080, *p* = 0.007; **Fig. 5A**) but not WD rats (*p* = 0.499). GE2 treatment reduced cytokine levels in females (F_(1,26)_ = 6.050, R^2^ = 0.189, *p* = 0.005; Sex and Treatment interaction: F_(1,54)_ = 3.763, R^2^ = 0.057, *p* = 0.022), but not males (*p* = 0.414). The main effect of treatment was also significant (F_(1,54)_ = 4.949, R^2^ = 0.075, *p* = 0.008) but there were no other significant main effects or interactions p ≥ 0.194).

**Figure 5.**
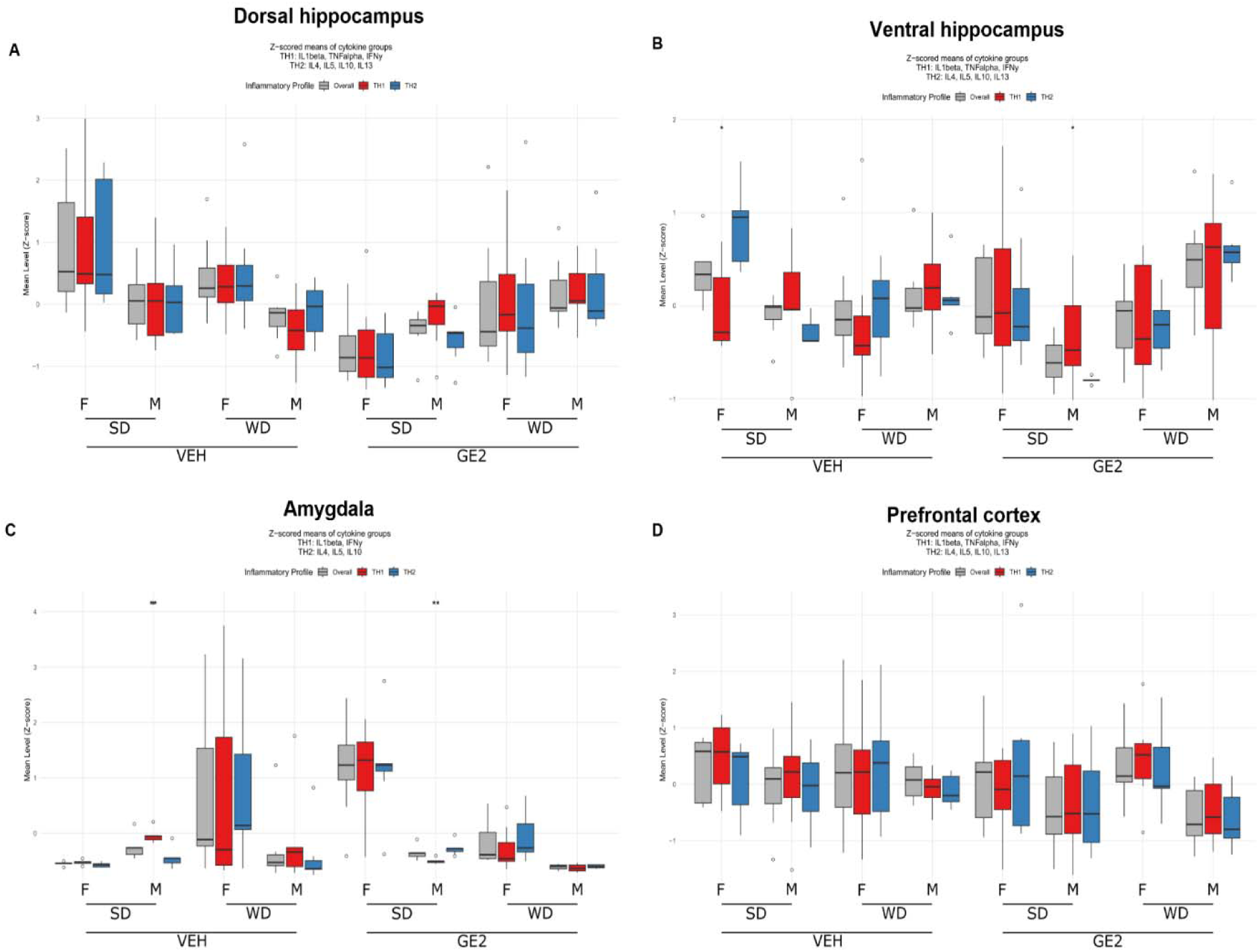
Effects of sex, diet and GE2 treatment on cytokine levels in the dorsal hippocampus, ventral hippocampus, amygdala and prefrontal cortex. Z-score composites for cytokines in the dorsal hippocampus (A) ventral hippocampus (B), amygdala (C) and prefrontal cortex (D). For Z-score composites the total cytokine score is presented in grey, TH1-produced cytokines in red and TH2-produced cytokines in blue. ** p < 0.01, *** p < 0.001, **** p < 0.0001 and # p < 0.100. n = 6-8 (VEH) and 6-8 (GE2) per diet and sex. SD = standard diet; WD = western diet; VEH = vehicle; GE2 = GLP-1-E2; F = female, M = male. Data are presented as mean ± confidence intervals (CIs). 0.245).

To better characterize the immunological profile associated with treatment, we examined cytokines commonly associated with T helper (TH) cell polarization, specifically TH1- and TH2-related cytokines. TH1 and TH2 refer to functional subsets of CD4+ T cells—typically associated with the production of IL-1β, IFN-γ, and TNFα (TH1), and IL-4, IL-5, IL-10, and IL-13 (TH2). However, it is important to note that these cytokines can also be produced by other cell types, including microglia and macrophages, particularly in the brain. Thus, while the TH1/TH2 classification provides a useful framework for interpreting cytokine profiles, the present study does not directly assess T cell populations, and the observed changes in cytokine levels may reflect activity from a broader range of immune and glial cell types.

We observed a reduction TH1-related cytokines after GE2 treatment in females (*p* = 0.016), but not in males (*p* = 0.474; sex and treatment interaction: F_(1,54)_ = 4.859, R^2^ = 0.076, *p* = 0.013). Furthermore, we observed a reduction in TH1-related cytokines after GE2 treatment in SD-fed rats (F_(1,25)_ = 5.527, R^2^ = 0.101, *p* = 0.010; diet and treatment interaction: F_(1,54)_ = 5.709, R^2^ = 0.089, *p* = 0.008), but not WD (*p* = 0.524). We found no other significant main effects or interactions (*p* ≥ 0.117).

A reduction in TH2-related cytokines was also observed after GE2 treatment in females (*p* = 0.005; sex and treatment interaction: F_(1,54)_ = 4.518, R^2^ = 0.065, *p* = 0.026), but not males (*p* = 0.654). Reduced TH2-related cytokine levels were present in GE2-treated SD rats (F_(1,25)_ = 14.580, R^2^ = 0.368, *p* < 0.001; diet and treatment interaction: F_(1,54)_ = 5.824, R^2^ = 0.084, *p* = 0.010), but not WD-fed rats (*p* = 0.945). The main effect of diet was also significant (main effect of diet: F_(1,54)_ = 6.413, R^2^ = 0.093, *p* = 0.008), and there was a trend towards a significant main effect of treatment (main effect of treatment: F_(1,54)_ = 2.791, R^2^ = 0.040, *p* = 0.080). There were no other significant main effects or interactions (*p* ≥ 0.324). Results for individual cytokines can be found in the Supplement.

#### 3.8.2. Ventral hippocampus

In this area we observed that WD increased cytokine levels in males only, and GE2 treatment reduced cytokine levels in SD-fed males and females (Fig. 5B).

WD increased cytokine levels in males compared to SD (F_(1,12)_ = 3.172, R^2^ = 0.209, *p* = 0.010). However, in females, SD-fed rats had higher cytokine levels than WD-fed rats (F_(1,11)_ = 3.705, R^2^ = 0.252, *p* = 0.007), this was primarily driven by high levels of TH2-produced, or anti-inflammatory, cytokines (F_(1,11)_ = 8.701, R^2^ = 0.442, *p* = 0.002). SD-fed females also had higher cytokine levels than SD-fed males (F_(1,10)_ = 8.501, R^2^ = 0.460, *p* = 0.001).

GE2 treatment significantly reduced cytokine levels in SD-fed females (F_(1,11)_ = 4.936, R^2^ = 0.310, *p* < 0.001; sex, diet and treatment interaction: F_(1,54)_ = 3.844, R^2^ = 0.051, *p* = 0.002), but not in WD-fed females (*p* = 0.781). In males, GE2 treatment reduced cytokine levels in SD-fed rats (F_(1,12)_ = 5.740, R^2^ = 0.324, *p* < 0.001), contrarily there was a trend towards increased cytokine levels after GE2 treatment in WD-fed rats (F_(1,12)_ = 2.192, R^2^ = 0.154, *p* = 0.073). We also found a significant interaction between diet and treatment (F_(1,54)_ = 5.225, R^2^ = 0.080, *p* = 0.007) and the main effect of treatment also significant (F_(1,54)_ = 4.949, R^2^ = 0.075, *p* = 0.008). No other significant main effects or interactions (*p* ≥ 0.194).

Assessing TH1-related cytokines individually, we found a significant main effect of diet, with increasing levels in SD-fed subjects compared to WD-fed subjects (F_(1,54)_ = 4.805, R^2^ = 0.079, *p* = 0.008). We found higher levels of TH2-related cytokines in SD-fed females compared to WD-fed females (F_(1,11)_ = 8.701, R^2^ = 0.442, *p* = 0.002). Furthermore, GE2 treatment reduced the levels of TH2-related cytokines in SD-fed rats of both sexes (females: F_(1,11)_ = 9.253, R^2^ = 0.457, *p* < 0.001; males: F_(1,12)_ = 51.496, R^2^ = 0.811, *p* < 0.001; sex, diet, treatment interaction: F_(1,54)_ = 9.078, R^2^ = 0.081, *p* < 0.001). This effect was not present in WD-fed females (*p* = 0.474), and an opposite effect was seen in WD-fed males, in which GE2 treatment increased the levels of TH2-produced cytokines (F_(1,12)_ = 6.833, R^2^ = 0.363, *p* = 0.003). There was also a significant interaction between diet and treatment (F_(1,54)_ = 12.950, R^2^ = 0.116, *p* < 0.001), sex and treatment (F_(1,54)_ = 6.074, R^2^ = 0.054, *p* < 0.001), sex and diet (F_(1,54)_ = 14.022, R^2^ = 0.125, *p* < 0.001), as well as significant main effects of treatment (F_(1,54)_ = 6.377, R^2^ = 0.057, *p* < 0.001), diet (F_(1,54)_ = 7.918, R^2^ = 0.071, *p* < 0.001) and sex (F_(1,54)_ = 7.558, R^2^ = 0.068, *p* < 0.001). Results for individual cytokines can be found in the Supplement.

#### 3.8.3. Amygdala

In the amygdala we observed elevated cytokine levels in response to a WD in females only and GE2 treatment reduced cytokine levels in males regardless of diet, but increased it in SD-fed females only (**Fig. 5C**). WD increased the cytokine levels in the amygdala compared to SD in females (F_(1,51)_ = 3.958, R^2^ = 0.265, *p* = 0.042; sex, diet and treatment interaction: F_(1,54)_ = 7.570, R^2^ = 0.084, *p* = 0.002), but not in males (*p* = 0.849).

GE2 reduced cytokine levels in the amygdala of WD-fed males (F_(1,12)_ = 2.045, R^2^ = 0.146, *p* = 0.018), but not WD-fed females (*p* = 0.157). GE2 also reduced cytokine levels in SD-fed males (F_(1,12)_ = 10.971, R^2^ = 0.478, *p* < 0.001), but increased cytokine levels in SD-fed females (F_(1,11)_ = 11.371, R^2^ = 0.508, *p* = 0.003). There were trends for interactions between diet and treatment (F_(1,54)_ = 10.746, R^2^ = 0.036, *p* = 0.053), sex and treatment (F_(1,54)_ = 3.217, R^2^ = 0.033, *p* = 0.053) and sex and diet (F_(1,54)_ = 2.970, R^2^ = 0.033, *p* = 0.065). The main effect of sex was significant (F_(1,54)_ = 13.770, R^2^ = 153, *p* < 0.001) and there was a trend towards a significant main effect of diet (F_(1,54)_ = 2.623, R^2^ = 0.029, *p* = 0.086), but no significant main effect of treatment (*p* =

GE2 treatment decreased the levels of TH1- and TH2-related cytokines in SD males (TH1: F_(1,12)_ = 77.656, R^2^ = 0.866, *p* < 0.001; TH2: F_(1,12)_ = 13.013, R^2^ = 0.520, *p* < 0.001). However, we found an increase in TH1- and TH2-related cytokines in SD females (TH1: F_(1,11)_ = 13.075, R^2^ = 0.543, *p* = 0.004; sex, diet and treatment interaction: F_(1,54)_ = 7.970, R^2^ = 0.100, *p* = 0.004; TH2: F_(1,11)_ = 15.013, R^2^ = 0.577, *p* = 0.004; sex, diet and treatment interaction: F_(1,54)_ = 8.161, R^2^ = 0.077, *p* < 0.001). A reduction in TH1-related cytokines was also present in vehicle-treated WD-fed males compared to SD-fed males (F_(1,12)_ = 2.424, R^2^ = 0.168, *p* = 0.038), but this effect was not present in females (*p* = 0.188), and no significant differences in TH2-related cytokines were present between treatment groups in WD-fed subjects (*p* ≥ 0.115).

For TH1-related cytokines, we also found a significant interaction between diet and treatment (diet and treatment interaction: F_(1,54)_ = 8.136, R^2^ = 0.102, *p* = 0.003) and the main effect of sex was also significant (main effect of sex: F_(1,54)_ = 9.488, R^2^ = 0.119, *p* = 0.001). For TH2-related cytokines, we also found a significant interaction between diet and treatment (diet and treatment interaction: F_(1,54)_ = 13.881, R^2^ = 0.130, *p* = 0.002), sex and treatment (sex and treatment interaction: F_(1,54)_ = 4.068, R^2^ = 0.038, *p* = 0.028) and sex and diet (sex and diet interaction: F_(1,54)_ = 6.159, R^2^ = 0.058, *p* = 0.005). Furthermore, the main effect of sex (main effect of sex: F_(1,54)_ = 20.908, R^2^ = 0.196, *p* < 0.001) and diet (diet: F_(1,54)_ = 4.155, R^2^ = 0.196, *p* < 0.001) were significant.

The levels of IL-6, IL-13 and TNFα were below detection in all of the samples in the amygdala. Furthermore, 59% of IL-5 and 50% of IFNγ samples were below detection. Differences in the levels of these cytokines could therefore not be analyzed. Results for individual cytokines above detection threshold can be found in the Supplement.

#### 3.8.4. PFC

Females had significantly higher levels of cytokine levels in the PFC than males (main effect of sex: F_(1,74)_ = 3.597, *p* = 0.022, R² = 0.061; **Fig. 5D**). When assessing TH1- and TH2-related cytokines separately, we observed that females had higher levels of TH2-related cytokines than males (main effect of sex for TH2: F_(1,55)_ = 4.615, R^2^ = 0.083, *p* = 0.018); an effect not found for TH1-related cytokines, nor were there any other significant main effects or interactions present, although there was a trend towards a diet effect (main effect of diet: *p* = 0.060; all other *p* ≥ 0.117).

## 4. Discussion

The results from this study revealed sex differences in metabolic and central aging biomarkers, as well as effects of GE2 on improving associative memory and aging biomarkers, in middle-aged rodents, regardless of diet. WD increased weight gain and plasma leptin levels and reduced PSD95 in the amygdala in both sexes, whereas GE2 treatment induced weight loss, elevated plasma PYY levels, and improved associative learning in both sexes. Although 8-week WD exposure led to significant weight gain, it did not lead to robust changes in contextual fear, central cytokine levels, neurogenesis or synaptic plasticity; however, we found certain sex-specific conditions in which diet affected these factors. Furthermore, we found sex differences in fat distribution patterns, blood glucose regulation, neurogenesis, cytokine levels and PSD95 expression in response to treatment. For instance, on WD females accumulated more visceral fat, whereas males accumulated more subcutaneous fat. Blood glucose regulation also differed, with GE2 improving glucose levels only in females. Neurogenesis in the DG was reduced with WD only in males, and although GE2 increased neurogenesis in both sexes this depended on diet (SD in females and WD in males). WD consumption in middle-age increased cytokine levels in the amygdala of females and in the ventral hippocampus of males, whereas GE2 treatment reduced cytokine levels in the hippocampus and amygdala, depending on diet and sex. Our findings reveal sex-specific vulnerabilities to diet-induced changes and demonstrate beneficial effects of GE2 regardless of diet, supporting its therapeutic potential, which may be further elucidated in future studies using more vulnerable models of obesity and diabetes, such as ob/ob or STZ-treated rodents. Furthermore, these findings underscore the complex interplay between sex, diet, and GE2 efficacy in middle age, emphasizing the need for tailored therapeutic strategies.

### 4.1. Sex-specific increases in subcutaneous and visceral fat in response to the WD

The 2025 obesity report from The Lancet reported that the worldwide prevalence of obesity, and being overweight, is higher in females than in males, which reaches its peak around middle age (50 years) in both sexes (16). This is consistent with the weight-gain observed at middle age in this study in rodents, as females gained more weight after 4 weeks on a WD than their male counterparts. Females and males often also carry differential amounts of adipose tissue which is stored in a site-specific manner, with females carrying more subcutaneous fat, whereas males often have more visceral fat (77). This sex difference is mainly driven by levels of estrogens after menopause in which human females show a shift in their main fat distribution from subcutaneous to visceral, which more closely resemble the distribution in males (46). In the rodents in the current study, we observed that both sexes gained subcutaneous fat (IWAT) on a WD, but only females on the WD gained visceral fat (GWAT), consistent with changes in fat distribution at this age in humans as described above. Therefore, we may be capturing a susceptible period, in middle age, for weight gain in females, directed towards increased fat storage in their visceral depots.

Consistent with previous results, four weeks of GE2 treatment reduced body weight (63). GE2 treatment resulted in loss of visceral fat (GWAT), however, this was dependent on diet and sex. More specifically, GE2 reduced visceral fat in males on SD, and tended to reduce this fat depot in WD-fed females. Of note, SD females had low levels of visceral fat, whereas males on SD had similar levels to their WD-fed counterparts. Collectively, these findings highlight sex-specific effects of diet and GE2 treatment on fat distribution and weight loss during middle age which may influence the effectiveness of GE2 treatment in different metabolic contexts.

### 4.2. GE2 treatment reduced basal blood glucose levels in females, but not males. Furthermore, this dose was not sufficient to reduce blood glucose after a glucose challenge in either sex

Females are more insulin sensitive than males (79). In the current study, WD increased fasting glucose and impaired glucose tolerance in the GTT, but only in females. However, under SD conditions, males had higher fasting glucose levels than females, despite higher insulin levels, suggesting reduced insulin sensitivity or early insulin resistance in males. GE2 treatment reduced basal blood glucose in females but did not improve glucose tolerance in the GTT. These data indicate that females may be more sensitive to the blood-glucose regulating effects of the GE2 conjugate than males. This finding is inconsistent with previous results that showed a potent blood glucose regulating effect of GE2 in males, however the mice in previous study (65) were given dose of GE2 that was 6-20 times higher than the one used in the current study (63). Data collected in humans using GLP-1 receptor agonists do, however, indicate that females are more sensitive to the blood glucose regulating effects of these treatments, and report higher incidences of symptomatic hypoglycemia than males (80). Collectively these findings demonstrate that although GE2 treatment effectively reduces body weight in both sexes, females may be more sensitive to its blood-glucose regulating effects.

### 4.3. GE2 treatment increased plasma GLP-1, but only in females, whereas plasma PYY increased in both sexes

Consistent with previous studies (81,82), we found that WD led to increases in plasma leptin compared to SD in both sexes. Interestingly, increases in plasma GLP-1 in GE2-treated rats were observed only in females. Females have previously been demonstrated to have substantially higher GLP-1 levels postprandially (83,84), and males have higher levels for Dipeptidyl Peptidase-4 (DPP-4), the main metabolizer of GLP-1 (85). This sex difference may be the result of higher endogenous GLP-1 remaining after their last meal or prolonged exposure following treatment due to differences in meal-stimulated GLP-1 secretion and GLP-1 metabolism. Of note, treatment and last meal occurred at the same time in both males and females. Furthermore, E2 increases GLP-1 secretion, and GLP-1 levels decrease in response to ovariectomy, an effect which can be rescued by E2 administration (86). It is therefore possible that GE2 increased GLP-1 secretion in females at middle age; a mechanism which is not present in males. We also found that males had higher levels of plasma insulin than females. Females are more sensitive to insulin and males are more vulnerable to insulin resistance (87), which may explain this sex difference.

GE2 treatment also led to increased levels of plasma PYY, an appetite-suppressing hormone produced in the small intestine (88), in both sexes. Levels of PYY are body-weight dependent, with normal weight subjects presenting with higher levels of plasma PYY than obese humans, and elevations in PYY have also been seen after weight loss, especially in combination with exercise (89). Levels of PYY were increased after GE2 treatment regardless of diet or sex which is in line with the weight reductions seen in these rats.

### 4.4. GE2 improved associative memory in both sexes

Previous findings demonstrated that GLP-1 improves hippocampus-dependent learning and memory in both standard-(34) and high-fat diet fed male rodents (35). A recent meta-analysis found that E2 positively affects spatial memory in the Morris water maze in females, and E2 can potentiate the actions of GLP-1 on other centrally mediated behaviors, such as food intake and reward (61,62). Here we found increased contextual and cued fear memory in both sexes after GE2 treatment.

Surprisingly, WD reduced cued fear memory only in middle-aged males, but did not significantly influence contextual fear conditioning in either sex, which is consistent with a study assessing the impact of high-fat diet in middle-aged male and female mice on the fear conditioning task reported that only males were affected by the diet (90). High-fat or WD has equivocal effects on hippocampus-based memory tasks (91–98), however in tasks using amygdala-based memory task, male mice were found to have selectively impaired cued fear memory in response to diet exposure (99), consistent with our results, or finding no significant impact of WD on hippocampus-dependent memory tasks (97,98). Of note, most of these studies were conducted exclusively in younger adult male rodents. Collectively, this indicates that males may be more vulnerable to the detrimental effects of high-fat or WD in middle age, especially on amygdala-dependent associative conditioning.

### 4.5. WD increased cytokine levels in the amygdala and ventral hippocampus in a sex-specific manner and GE2 reduces cytokine levels in the hippocampus and amygdala dependent on diet and sex, respectively

In the present study, we found sex- and site-specific increases in cytokine levels in response to WD exposure, in which males presented with elevated cytokine levels in the ventral hippocampus, whereas females presented with increased cytokine levels in the amygdala. These findings in the hippocampus are consistent with previous results in aged mice (8.5 months) that detected increased cytokine genes in response to a high-fat diet in males (100). Furthermore, one previous study conducted exclusively in females demonstrated that the amygdala, but not hippocampus, may be especially vulnerable to diet-induced changes in females, resulting in inflammatory dysregulation (101).

Treatment with GE2 was effective at reducing cytokine levels in the dorsal and ventral hippocampus, in rats fed a SD, but not WD. Furthermore, GE2 reduced cytokine levels in the amygdala, regardless of diet, but only in males. The cytokine-level reductions in these areas were found to encompass both TH1 and TH2 produced cytokines, using this classification as a framework for pro- and anti-inflammatory signaling rather than implying specific cellular sources. Previous studies show that GLP-1 receptor agonist treatment reduced microglial activation and reversed lipopolysaccharide (LPS)-induced cytokine-level increases in the hippocampus (38,102,103), indicating that these properties are preserved in its conjugated form. Surprisingly, GE2 treatment in SD females resulted in increased cytokine levels in the amygdala, an effect which was not observed in other groups or brain regions. The females in this group had low body weights and did not lose weight during treatment, whereas middle-aged males on the same diet had higher percentual adipose tissue and lost weight after GE2 treatment. It could therefore be possible that the treatment is not as beneficial to this group and may instead be detrimental. However, this effect appears to be centralized to the amygdala and future studies are needed to determine whether this region-specific cytokine increase is negative or positive. Collectively, GE2 reduced cytokine levels in both the dorsal and ventral hippocampus, dependent on diet and sex; GE2 also reduced amygdala cytokines in males, regardless of diet, but increased them in SD-fed females.

### 4.6. WD reduced neurogenesis in the DG in WD-fed males and GE2 treatment restored neurogenesis in males. GE2 increased neurogenesis in the DG in SD-fed females

In the present study, GE2 treatment increased neurogenesis in the DG of middle-aged females fed a SD, consistent with a study examining effects of a GLP-1 receptor agonist treatment alone in middle-aged female mice (104). In males, the WD reduced neurogenesis in the DG, was commensurate with impaired cued memory after WD consumption in males only, consistent with previous studies in young adult and middle-aged males (105–107). WD-reduced neurogenesis in the DG was normalized by GE2 treatment, consistent with effects mediated by GLP-1 receptor agonist treatment in young adult male rodents (108–112) and adult and aged female mice (104). Collectively, GE2-treatment can increase neurogenesis in middle age, but this depended on diet and sex.

In the present study, we found that WD tended to reduce PSD95 in the amygdala only. This is in partial contrast to a study that found an increase in PSD95 in the hippocampus of male mice after GLP-1 receptor agonist treatment (113) in 5XFAD mice (a mouse model of Alzheimer’s disease), but not young, male wild-type young male mice.

### 4.7. Limitations

Although our study demonstrated clear improvements in body weight with WD, we did not observe strong effects of the diet on fear conditioning after 8 weeks of WD exposure in middle-aged rats and stronger effects may be observed with longer WD exposure or in disease models at risk for AD or diabetes. However, we did see clear sustained differences in body weight, fat and plasma leptin, indicating that the diet intervention was long enough to impact metabolic health. These metabolic changes may precede or contribute to inflammatory and cognitive alterations over a longer duration, suggesting that extended WD exposure or pre-existing metabolic vulnerability could amplify these effects. Future studies with extended diet exposure and/or the use of rodent disease models are necessary to further explore the potential benefits of GE2 treatment.

In this study we selected a low dose of GE2, based on prior research in rats (64) and the original study using this conjugate (63), after guidance from the Novo Nordisk team (BF, JD, BY). This decision was influenced by the fact that we included female rats, which are more sensitive to the effects of GLP-1 receptor agonists (29). Research indicates that females report more adverse effects when treated with GLP-1-based therapies (114). Additionally, individuals with lower body mass, including females, are more likely to experience nausea and other gastrointestinal side effects from GLP-1 receptor agonists (115). Furthermore, because our study was conducted in normal aging rats, without diabetes or other pathological conditions such as Alzheimer’s disease, we aimed to use the lowest effective dose to minimize potential adverse effects while still achieving beneficial effects on the brain. However, since some effects, such as those on cytokine levels, were only observed in SD-fed subjects, it is possible that the dose was not high enough to counteract the effects of the WD, and a higher dose may have led to reductions in WD-fed rats.

### 4.8. Conclusions

In conclusion, the findings from this study underscore the potential of GE2 treatment as a promising therapeutic approach for reducing body weight, visceral fat, and cytokine levels in the dorsal and ventral hippocampus, as well as the amygdala, depending on sex and diet, while increasing neurogenesis and associative memory. The observed sex-specific effects highlight the necessity of designing treatments that account for sex differences in neurodegenerative and metabolic diseases, considering that females are disproportionately affected by both Alzheimer’s disease and obesity. Although the WD did not induce robust neuroinflammatory changes in this model, future studies are warranted to investigate the impact of longer or more severe metabolic challenges on central cytokine levels and cognitive outcomes, as existing literature shows mixed findings. Moreover, the beneficial effects of GE2 on neurogenesis, cytokine levels, and memory suggest that this treatment may be particularly advantageous in populations already known to exhibit impairments in these domains, such as individuals with Alzheimer’s disease. As obesity, diabetes and dementia rates continue to rise globally, the intersection between metabolic dysfunction and cognitive decline emphasizes the urgent need for tailored, sex-specific therapies to address these overlapping health crises, offering hope for improving the quality of life for individuals at risk.

## Author contributions

**Jennifer E Richard** - Writing – original draft, review & editing, funding acquisition, Formal Analysis, Visualization; **Ahmad Mohammad** – Investigation; **Kimberly A Go** – Methodology, Project Administration; **Andrew J McGovern** – Formal Analysis **Rebecca K Rechlin** – Data Curation, Investigation; **Tallinn FL Splinter** – Data Curation, Investigation; **Stephanie E Lieblich** – Project Administration, Methodology; **Lara K Radovic** – Investigation; **Lydia Feng** – Investigation; **Samantha A Blankers** – Methodology, Visualisation; **Bin Yang** – Resources, Methodology; **Jonathan D Douros** – Resources, Methodology; **Brian Finan** – Resources, Methodology; **Liisa AM Galea** - Funding acquisition, Project administration, Resources, Supervision, Validation, Writing – review & editing.

## Acknowledgements

We would like to extend our heartfelt thanks to Mel Cevizci for her invaluable assistance during the GTT. We appreciate the support of Amanda Namchuk, Travis Hodges, Bonnie Lee, and Natascha Diekow, who contributed to the experiment endpoints and tissue collections. A special thanks to Niki Shahraki, Caitlin Ambrose, Lindsay Notte, Cherise Kwok, Aya Chahrour, Ziyi Zhang, Peixin Shi, Valerie Cheng, and Lele Ma for their dedicated help with behavioral scoring.

## Data availability

Data will be made available on request.

