## Supplement for "Sex-Specific Metabolic and Central Effects of GLP-1–Estradiol Conjugate in Middle-Aged Rats on a Standard or Western Diet"

**Supplementary Figures**

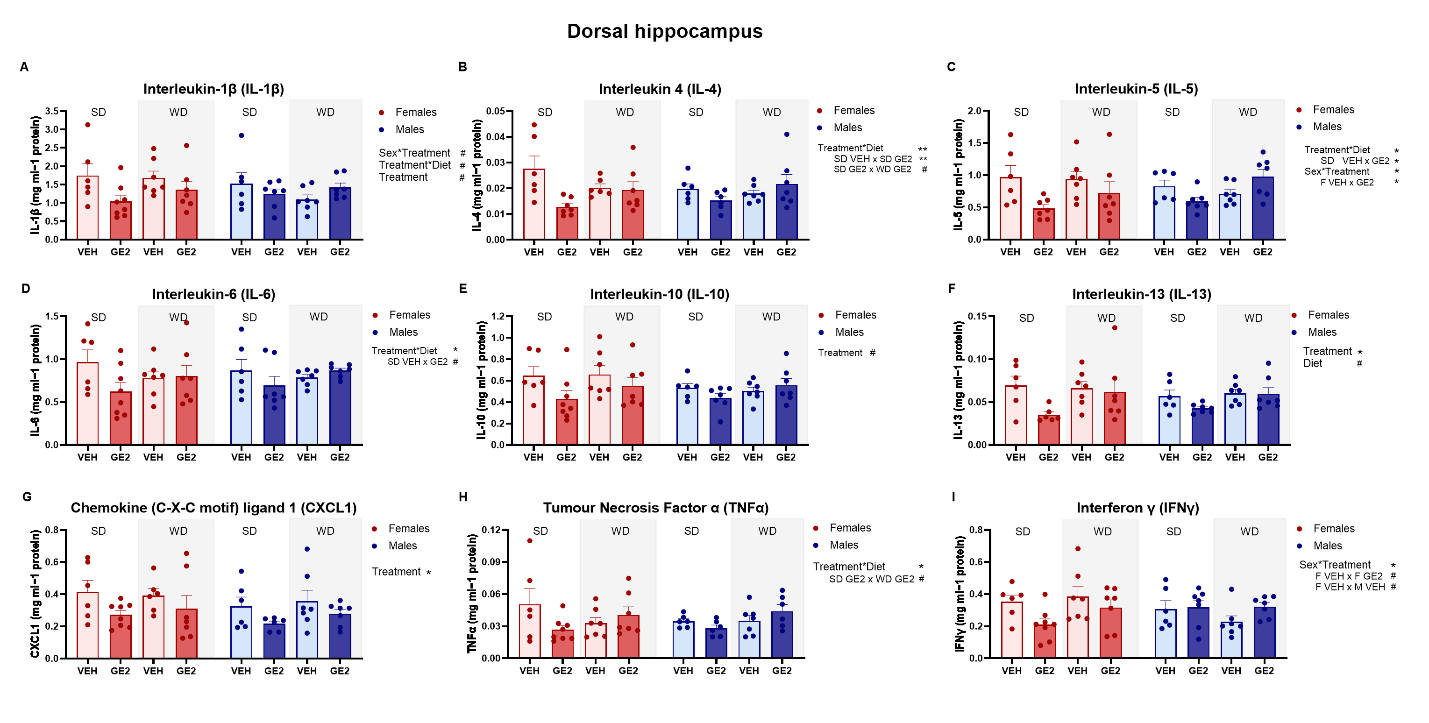

Supplementary Figure 1. **GE2 treatment reduced the levels of several cytokines in the dorsal hippocampus, with most prominent reductions in standard-diet fed females.** Levels of interleukin (IL)-1β (A), IL-4 (B), IL-5 (C), IL-6 (D), IL-10 (E), IL-13 (F), interferon (IFN) γ (G), C-X-C motif ligand 1 (CXCL1; H) and tumor necrosis factor (TNF) α (I). ** p < 0.01, *** p < 0.001, **** p < 0.0001 and # p < 0.100. n = 6-8 (VEH) and 6-8 (GE2) per diet and sex. SD = standard diet; WD = western diet, VEH = vehicle, GE2 = GLP-1-E2, F = female, M = male. Cytokine levels measured in relation to protein level and expressed as mg ml^-1^.

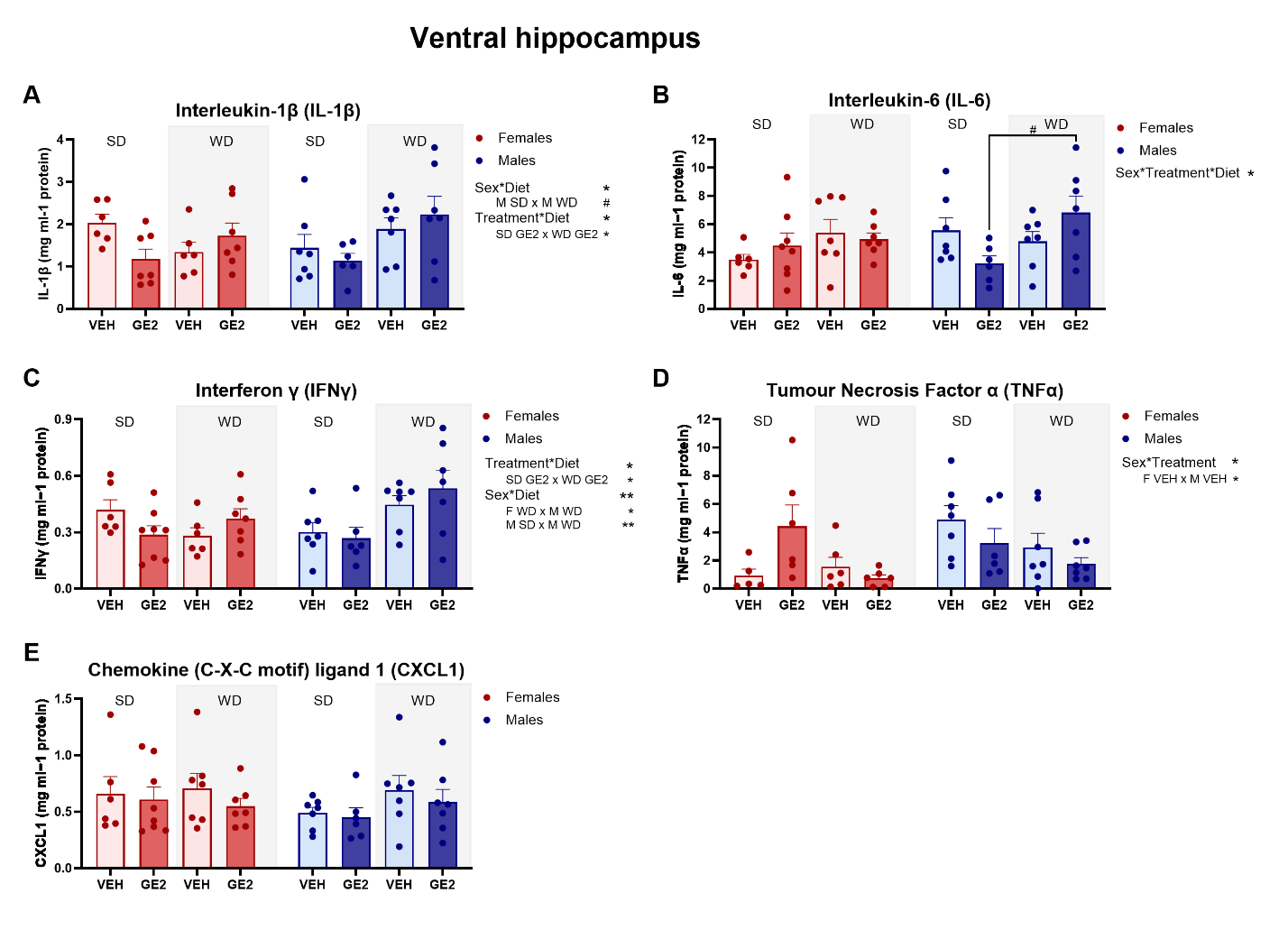

Supplementary Figure 2. **Cytokine levels in the ventral hippocampus.** Levels of IL-1β (A), IL-6 (B), IFNγ (C), TNFα (D) and CXCL1 (E) expressed as mg ml^-1^ protein. ** p < 0.01, *** p < 0.001 and # p <0.100. n = 5-8 (VEH) and 6-8 (GE2) per diet and sex. SD = standard diet; WD = western diet, F = female, M = male, VEH = vehicle, GE2 = GLP-1-E2.

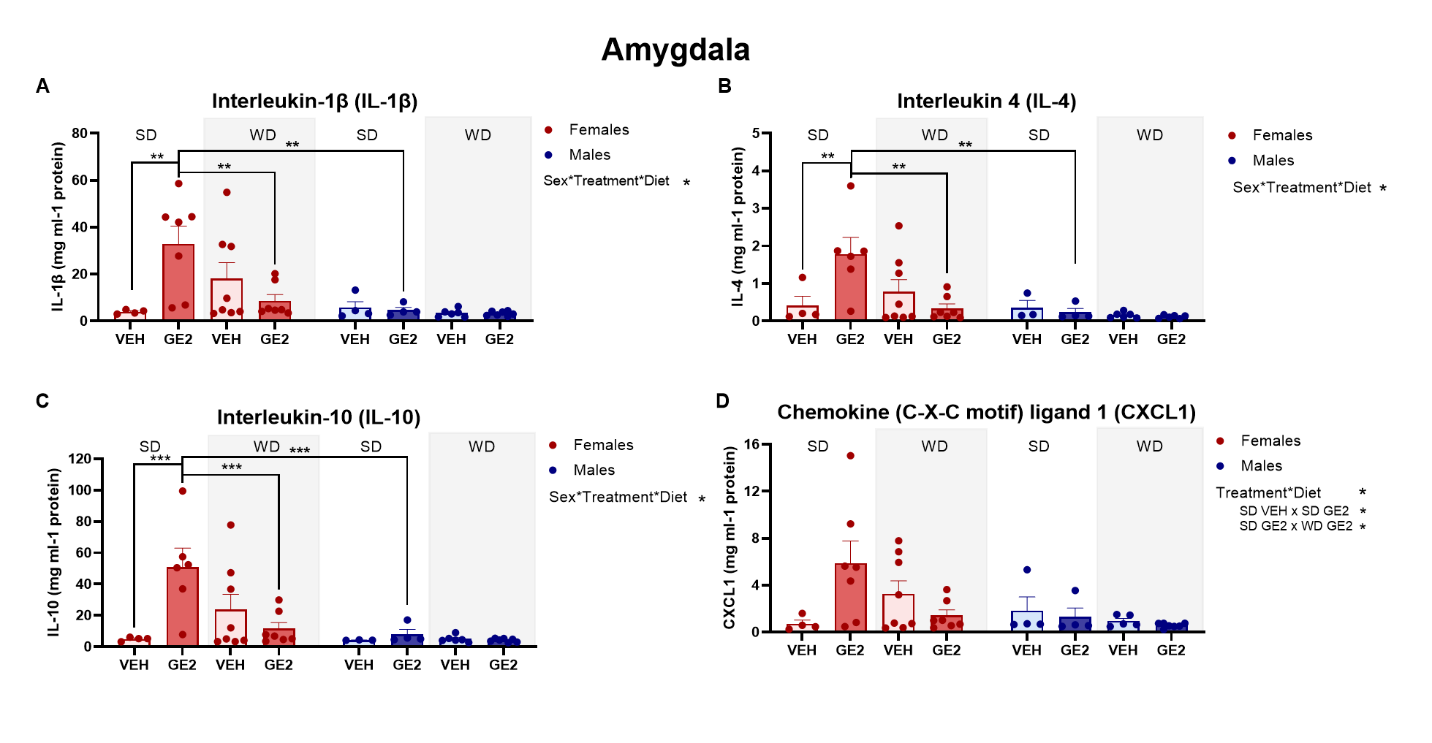

Supplementary Figure 3. **Cytokine levels in the Amygdala.** Levels of IL-1β (A), IL-4 (B), IL-10 (C), and CXCL1 (D) expressed as mg ml^-1^ protein. VEH = vehicle, GE2 = GLP-1-E2. * p < 0.05. n = 4-9 (VEH) and 4-8 (GE2) per diet and sex.

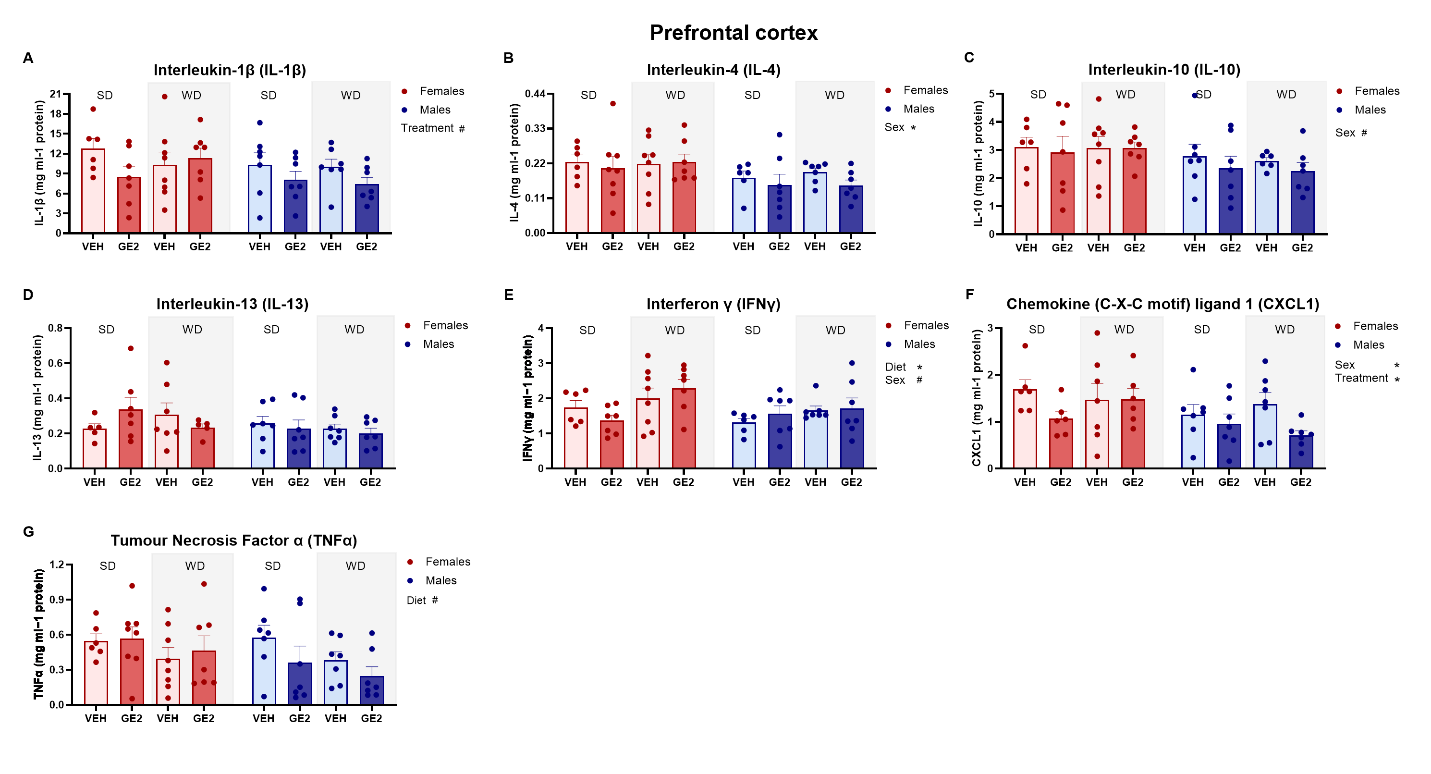

Supplementary Figure 4. **Cytokine levels in the prefrontal cortex (PFC).** Levels of IL-1β (A), IL-4 (B), IL-10 (C), IL-13 (D), IFNγ (E), CXCL1 (F) and TNFα (G) expressed as mg ml^-1^ protein. * p < 0.05, ** p < 0.01 and # p < 0.100. n = 5-8 (VEH) and 6-8 (GE2) per diet and sex. SD = standard diet; WD = western diet, F = female, M = male, VEH = vehicle, GE2 = GLP-1-E2.

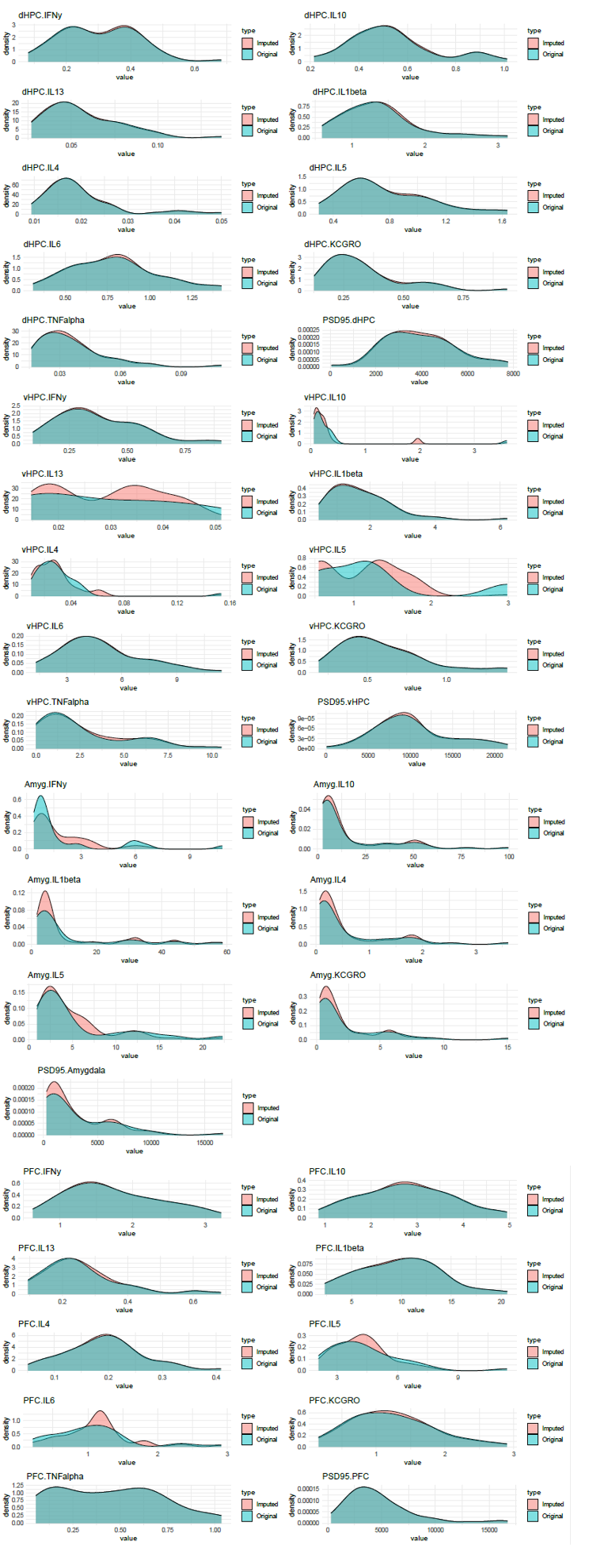

Supplementary Figure 5. Density Plots of Original and Imputed Cytokine Data. Distributions of original (teal) and imputed (pink) values for cytokines and PSD95 across four brain regions: dorsal hippocampus (dHPC), ventral hippocampus (vHPC), amygdala (Amyg), and prefrontal cortex (PFC), demonstrating the preservation of data structure following multiple imputation using the random forest method.

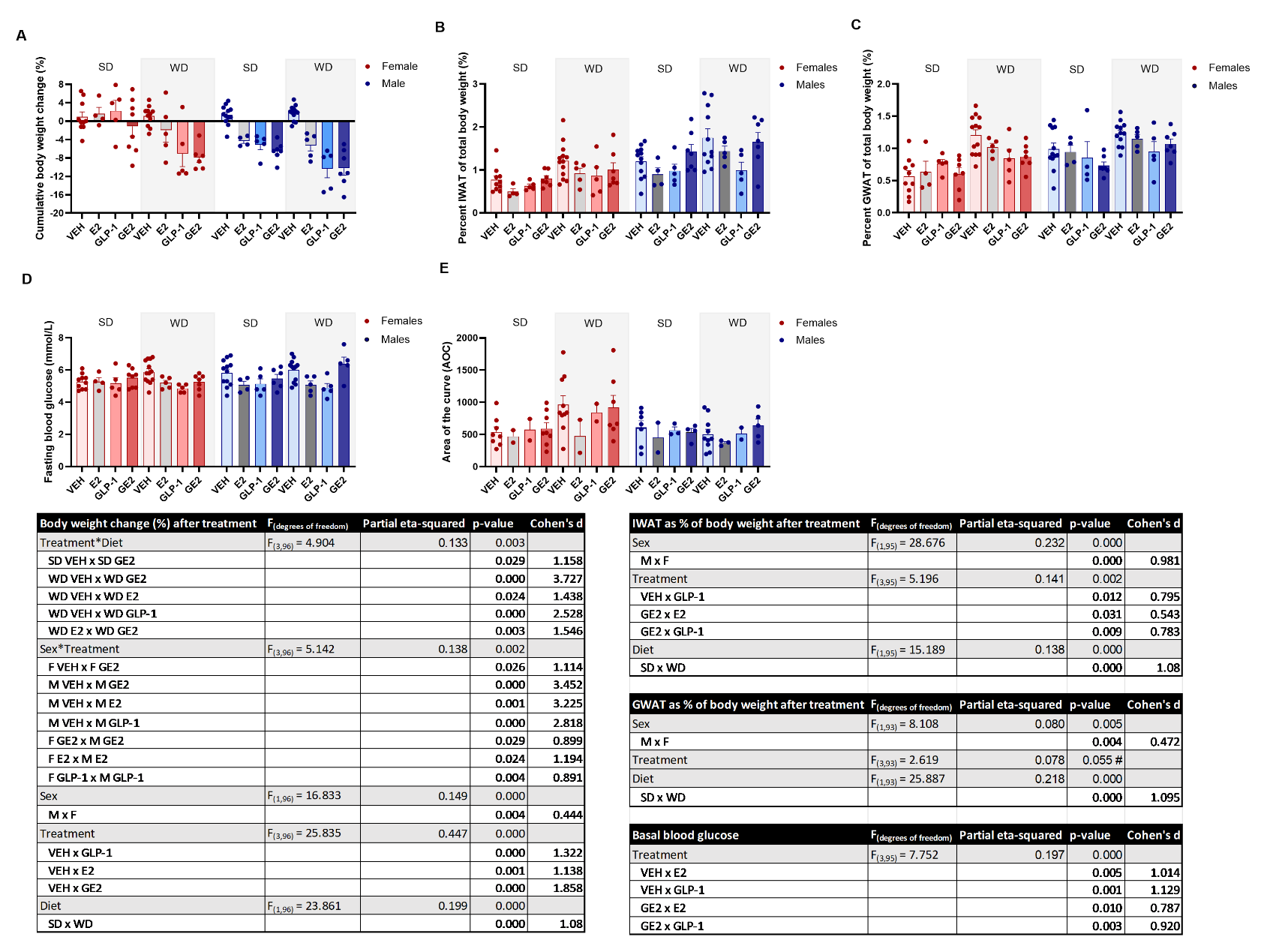

Supplementary Figure 6. **Body weight, fat and blood glucose for all groups.** Cumulative body weight change (A), percent inguinal white adipose tissue (IWAT; B), gonadal white adipose tissue (GWAT, C) basal blood glucose (D) and area under the curve for glucose tolerance test data (E) for vehicle, estradiol (E2), GLP-1 and GE2. Statistics for each graph are presented in the tables. One female (1 SD saline and one male (WD; GE2 treated) were lost during the experiment. Furthermore, unfortunately, due to an error in formulation, GTT testing failed in one cohort which resulted in fewer rodents in the GTT analysis for GLP-1 and E2 groups. n = 10-12 (vehicle), 4-5 (E2), 4-5 (GLP-1), 6-8 (GE2), per diet and sex for body weight and fat data and n = 8-11 (VEH), 2-3 (E2), 2-3 (GLP-1), 5-8 (GE2), per diet and sex for glucose-related measurements. F = females, M = males, SD = standard diet, WD = western diet, VEH = vehicle, GE2 = GLP-1-E2.

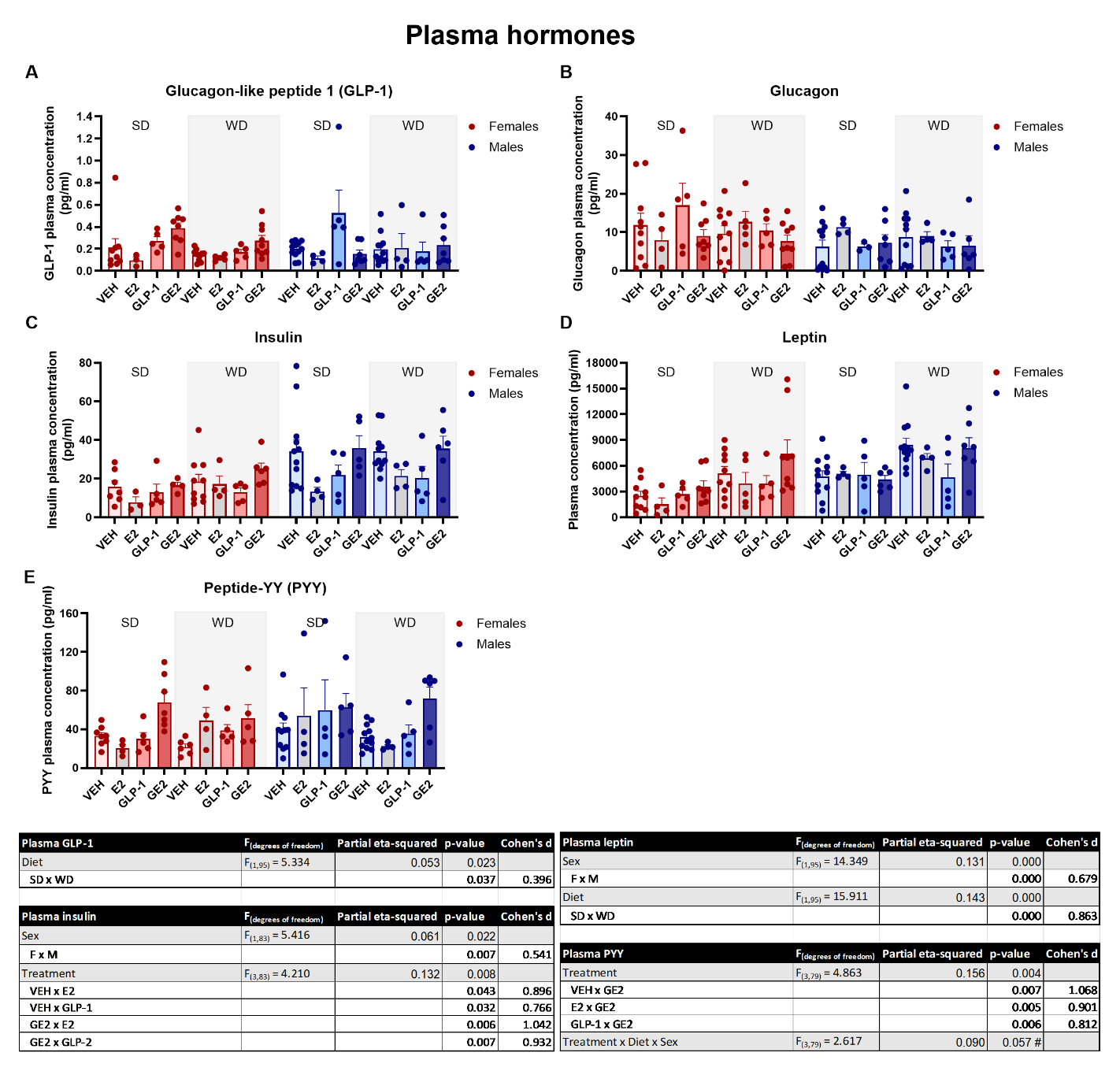

Supplementary Figure 7. **Levels of metabolic hormones in plasma for all groups.** Plasma levels of glucagon-like peptide-1 (GLP-1; A), glucagon (B), insulin (C), leptin (D), peptide YY (PYY; E) measured in pg/ml for vehicle, estradiol (E2), GLP-1 and GE2. Statistics for each graph are presented in the tables. n = 6-12 (VEH), 3-5 (E2), 3-5 (GLP-1), 4-9 (GE2), per diet and sex. F = female; M = male, SD = standard diet; WD = western diet; E2 = estradiol; GLP-1 = glucagon-like peptide-1, VEH = vehicle, GE2 = GLP-1-E2.

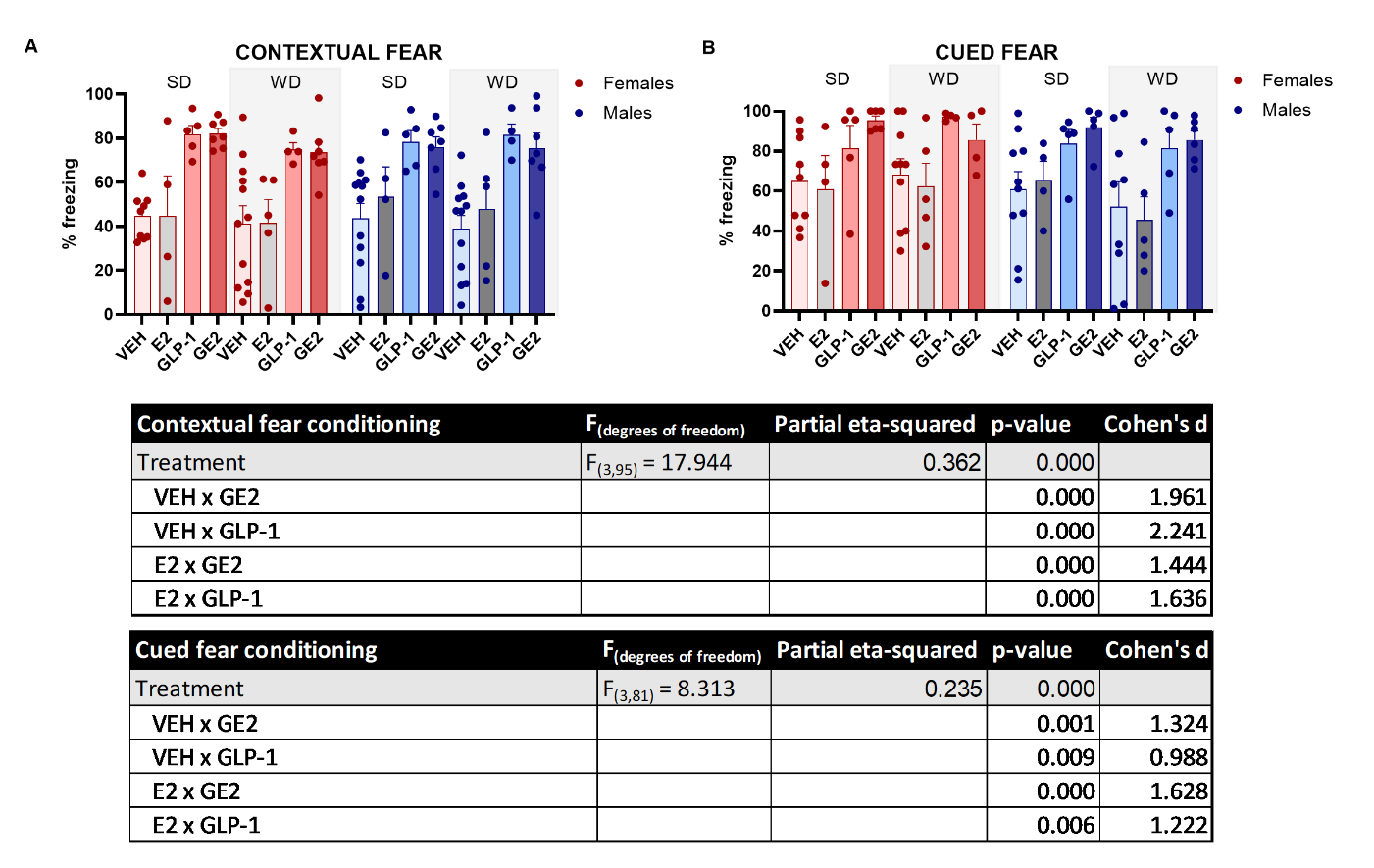

Supplementary Figure 8. **Fear conditioning test for all treatment groups.** Percent freezing in the contextual fear conditioning task (A) and the cued fear conditioning task (B). Four rats were excluded because they were out of frame and could not be scored (1 GLP-1 and 1 E2).Statistics for each graph are presented in the tables. n = 9-10 (VEH), 4-5 (E2), 4-5 (GLP-1), 4-6 (GE2), per diet and sex. F = female; M = male, SD = standard diet; WD = western diet; VEH = vehicle, E2 = estradiol; GLP-1 = glucagon-like peptide-1, GE2 = GLP-1-E2.

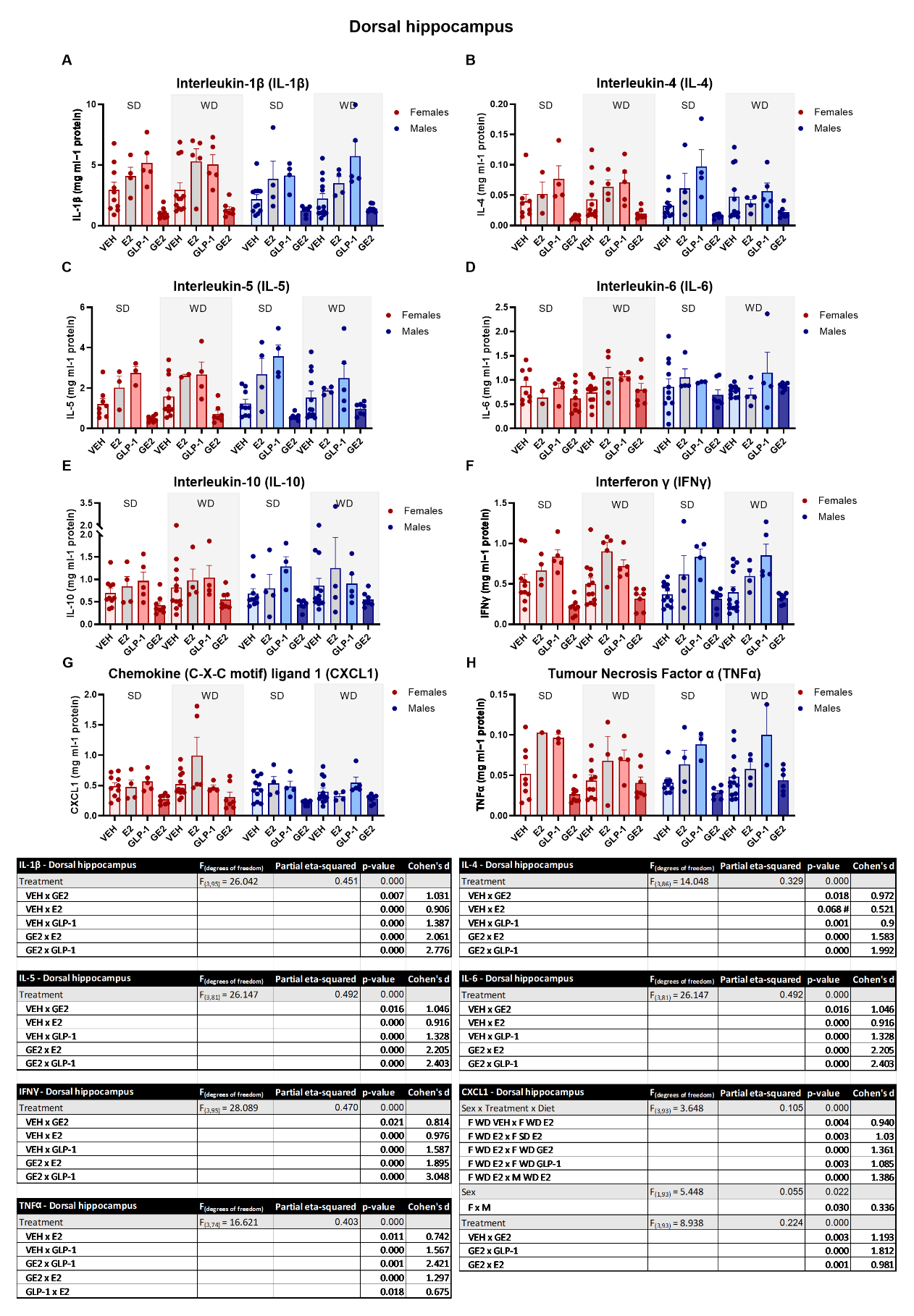

Supplementary Figure 9. **GE2 treatment reduced the levels of several cytokines in the dorsal hippocampus.** Levels of interleukin (IL)-1β (A), IL-4 (B), IL-5 (C), IL-6 (D), IL-10 (E), interferon (IFN) γ (F), C-X-C motif ligand 1 (CXCL1; G) and tumor necrosis factor (TNF) α (H). n = 8-13 (VEH), 1-5 (E2), 2-5 (GLP-1), 6-8 (GE2), per diet and sex. F = female; M = male, SD = standard diet; WD = western diet; VEH = vehicle, E2 = estradiol; GLP-1 = glucagon-like peptide-1, GE2 = GLP-1-E2.

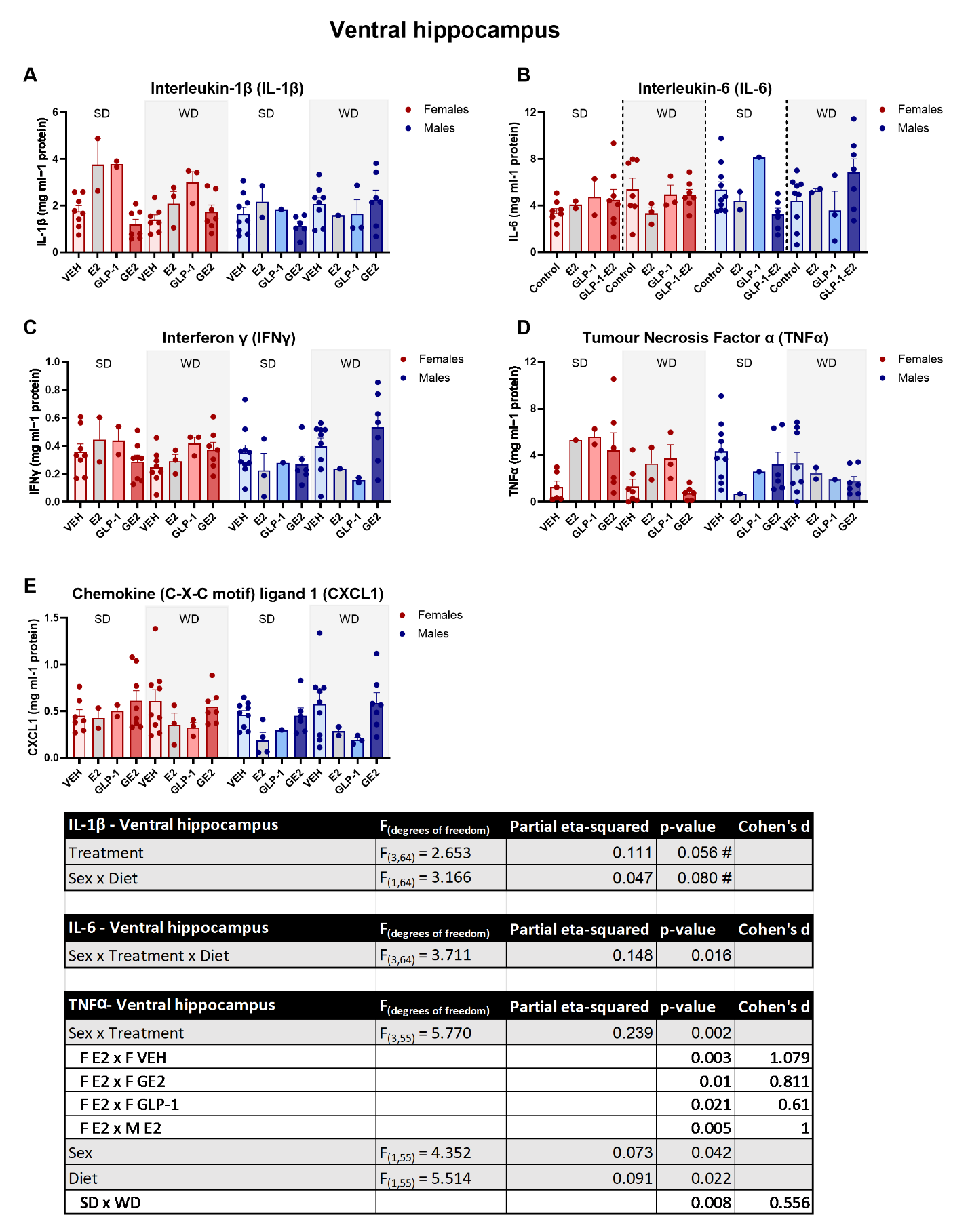

Supplementary Figure 10. **Few changes to cytokine levels in the ventral hippocampus.** Levels of interleukin (IL)-1β (A), IL-6 (B), interferon (IFN) γ (C), tumor necrosis factor (TNF) α (D) and C-X-C motif ligand 1 (CXCL1; E). n = 6-10 (VEH), 1-4 (E2), 1-3 (GLP-1), 6-10 (GE2), per diet and sex. F = female; M = male, SD = standard diet; WD = western diet; VEH = vehicle, E2 = estradiol; GLP-1 = glucagon-like peptide-1, GE2 = GLP-1-E2.

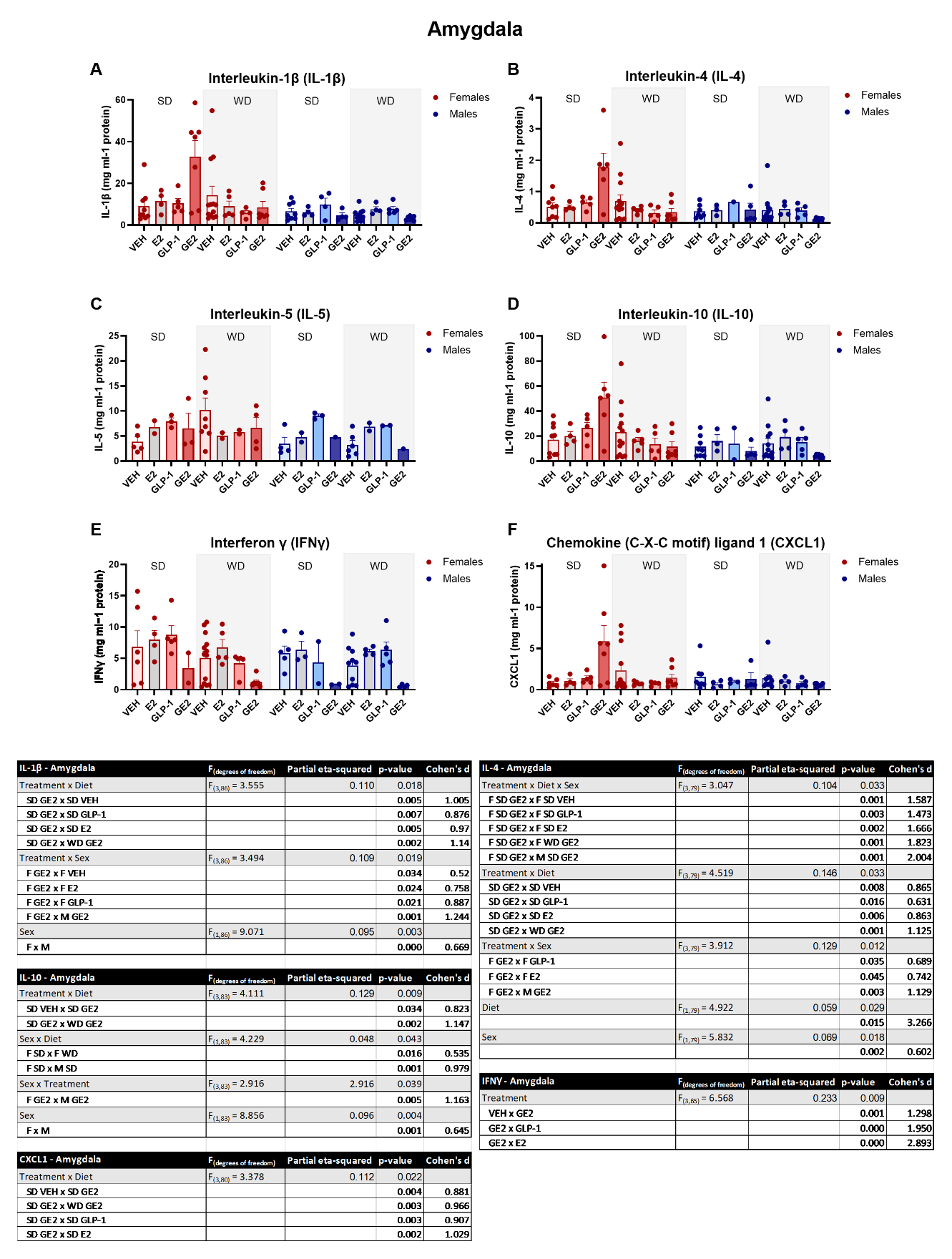

Supplementary Figure 11. **GE2 treated females on the standard diet had elevated levels of several cytokines.** Levels of interleukin (IL)-1β (A), IL-4 (B), IL-5 (C), IL-10 (D) interferon (IFN) γ (E) and C-X-C motif ligand 1 (CXCL1; F). n = 5-11 (VEH), 2-5 (E2), 1-5 (GLP-1), 1-5 (GE2), per diet and sex. F = female; M = male, SD = standard diet; WD = western diet; VEH = vehicle, GE2 = GLP-1-E2, E2 = estradiol; GLP-1 = glucagon-like peptide-1.

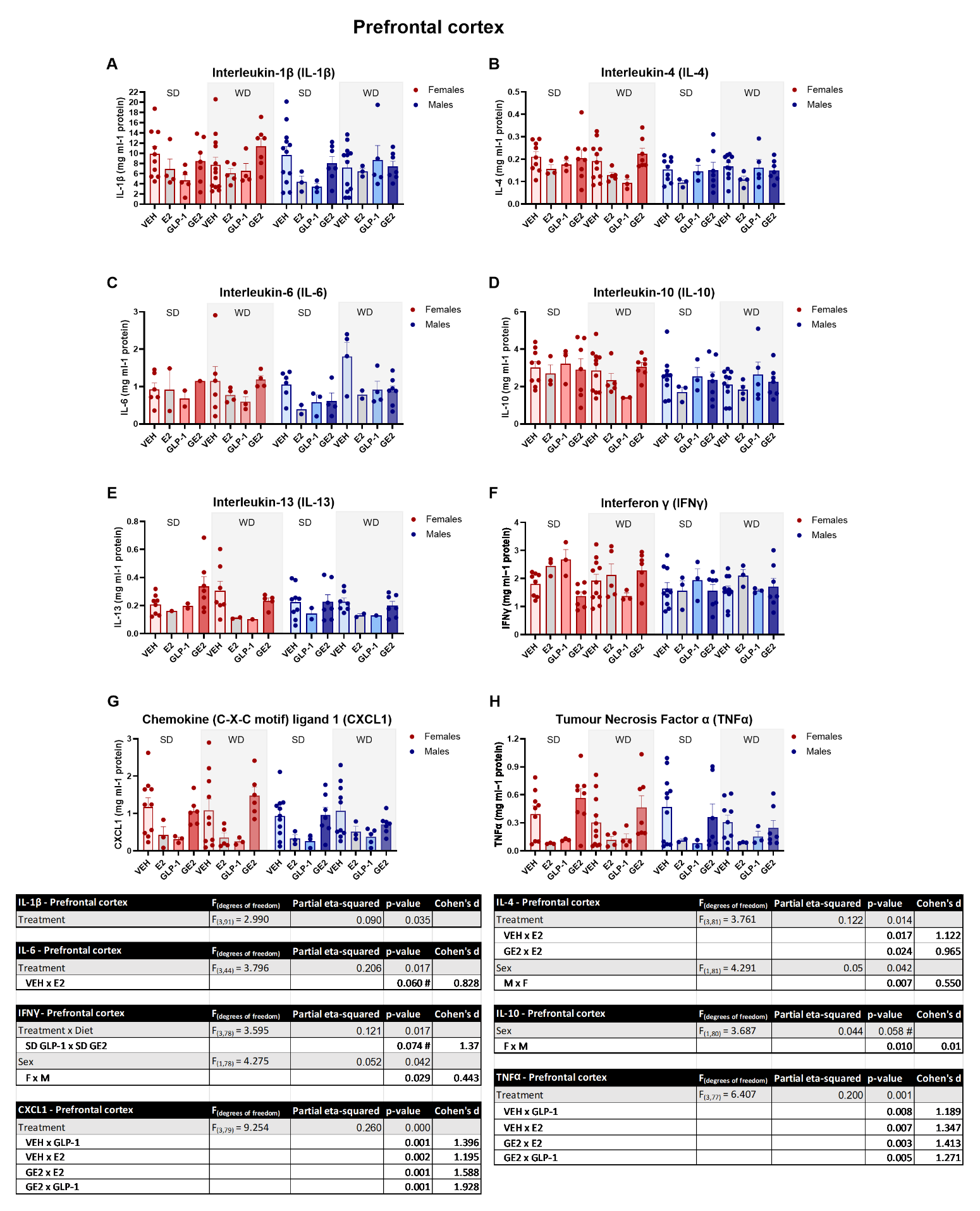

Supplementary Figure 12. **Cytokine levels in the prefrontal cortex.** Levels of interleukin (IL)-1β (A), IL-4 (B), IL-6 (C), IL-10 (D), IL-13 (E) interferon (IFN) γ (F), C-X-C motif ligand 1 (CXCL1; F) and tumor necrosis factor (TNF) α (G). n = 6-11 (VEH), 1-5 (E2), 1-5 (GLP-1), 1-8 (GE2), per diet and sex. F = female; M = male, SD = standard diet; WD = western diet; VEH = vehicle, E2 = estradiol; GLP-1 = glucagon-like peptide-1, GE2 = GLP-1-E2.

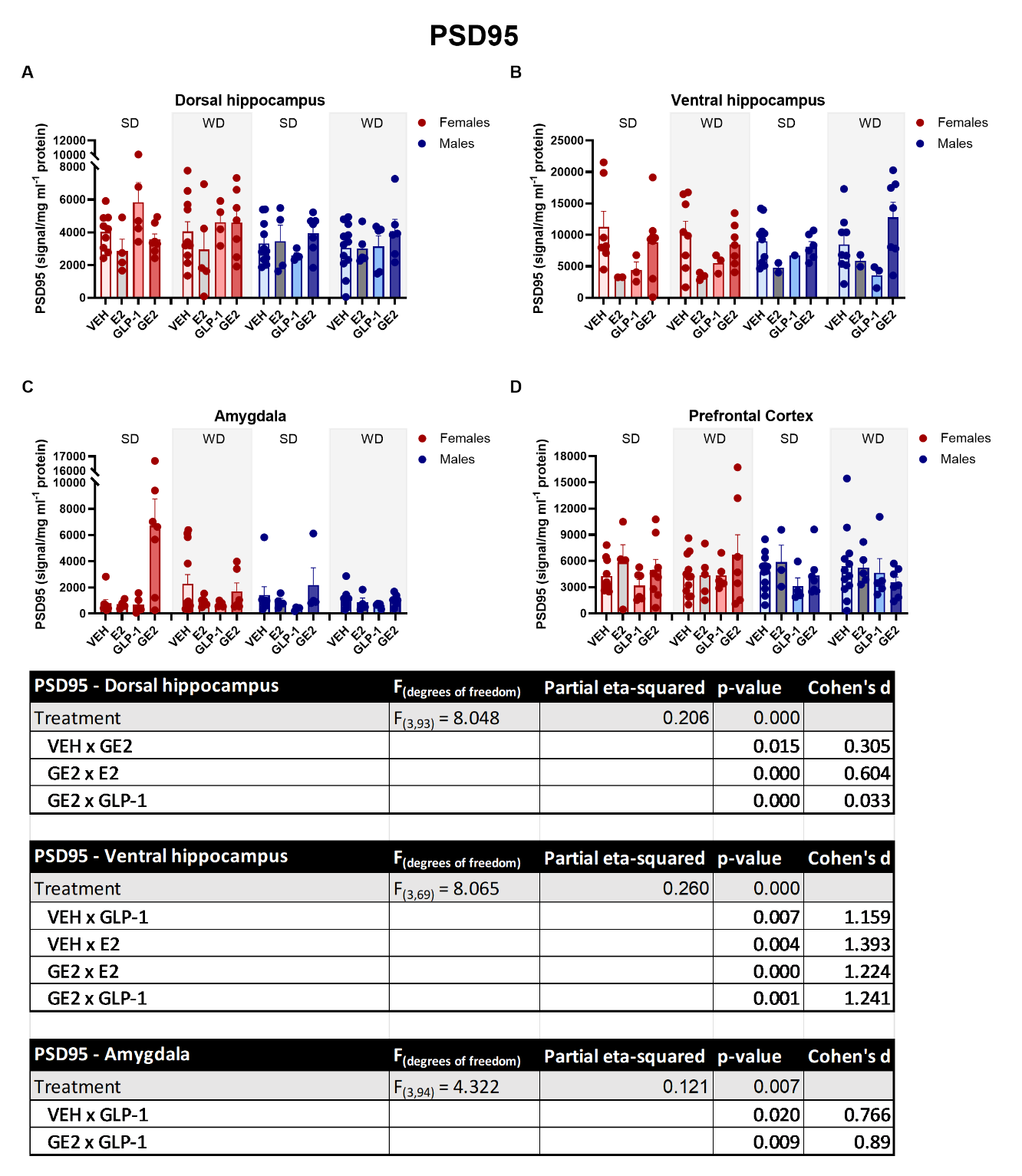

Supplementary Figure 13. **Levels of postsynaptic density protein 95 (PSD95) are increased in the dorsal hippocampus.** Levels of PSD95 in the dorsal hippocampus (A), ventral hippocampus (B), Amygdala (C), and prefrontal cortex (D). n = 8-13 (VEH), 1-5 (E2), 2-5 (GLP-1), 6-8 (GE2), per diet and sex. F = female; M = male, SD = standard diet; WD = western diet; VEH = vehicle, E2 = estradiol; GLP-1 = glucagon-like peptide-1, GE2 = GLP-1-E2.

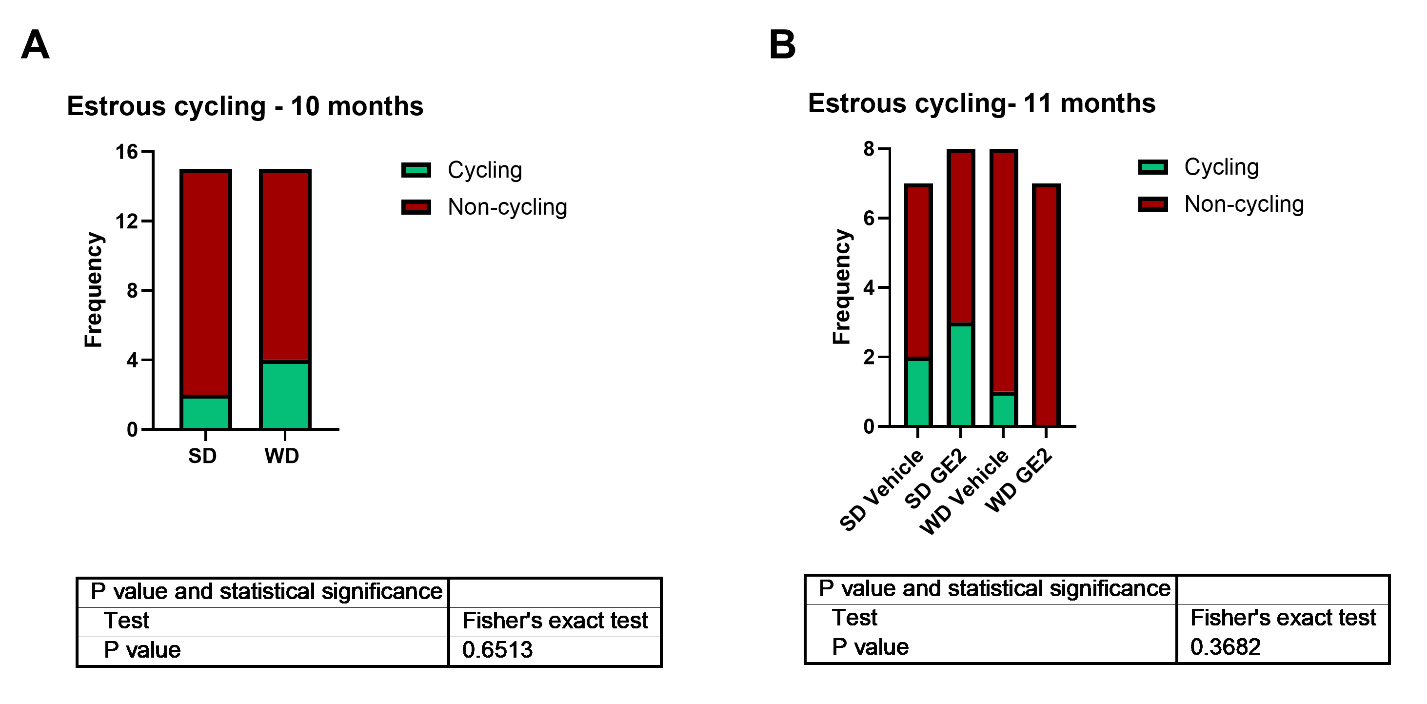

Supplementary Figure 14. Estrous cycling frequencies in females at 10 (A) and 11 months (B) of age. n = 15 per group (diet) for 10 months and n = 7-8 per group (diet and treatment) at 11 months. SD = Standard diet; WD = Western diet; GE2 = GLP-1-estradiol. Due to frequencies <1, the Fisher’s exact test was used to determine whether there was a non-random association between the categorical variables; however, neither test reached statistical significance, suggesting no strong evidence of association in our sample. Because very few females were still cycling at the time of behavioral testing we were, unfortunately, unable to assess the results based on cycling stays.

**Results - Individual cytokine levels after GLP-1-E2 and vehicle treatment**

*Dorsal hippocampus:* There was a trend towards lower levels of IL-1β in GE2-treated compared to vehicle-treated females (*p* = 0.054, Cohen’s *d* =0.910; treatment and sex interaction: F_(1,47)_ = 3.589, Ƞ_p_^2^ = 0.071, *p* = 0.064; **Supplementary Fig. 1A**), but not males (*p* = 0.858). There was also a trend towards lower levels of IL-1β specifically in SD-fed subjects (*p* = 0.068, Cohen’s *d* =0.849; treatment and diet interaction: F_(1,47)_ = 3.124, Ƞ_p_^2^ = 0.062, *p* = 0.084), but now WD (*p* = 0.995). There was also a trend towards a significant main effect of treatment (main effect of treatment: F_(1,47)_ = 3.095, Ƞ_p_^2^ = 0.062, *p* = 0.085), but no other significant main effects of interactions (*p* ≥ 0.342).

GE2 treatment significant reduced levels of IL-4 in SD subjects compared to vehicle (*p* = 0.005, Cohen’s *d* =1.336; treatment and diet interaction: F_(1,44)_ = 8.132, Ƞ_p_^2^ = 0.156, *p* = 0.007; **Supplementary Fig. 1B**), but not in WD-fed subjects (*p* = 0.598). In addition, levels of IL-5 were reduced in GE2- treated subjects fed SD, but not WD, compared to vehicle-treated subjects (*p* = 0.016, Cohen’s *d* =1.354; treatment and diet interaction: F_(1,46)_ = 5.646, Ƞ_p_^2^ = 0.109, *p* = 0.022; **Supplementary Fig. 1C**). Furthermore, GE2 treatment reduced IL-5 levels in females, regardless of diet (*p* = 0.020, Cohen’s *d* =0.980; treatment and sex interaction: F_(1,46)_ = 5.031, Ƞ_p_^2^ = 0.099, *p* = 0.030); an effect not seen in males (*p* = 0.863). There was also a trend towards a significant reduction in IL-6 in GE2, compared to vehicle treated, but only on the SD diet (*p* = 0.054, Cohen’s *d* =0.876; treatment and diet interaction: F_(1,47)_ = 4.860, Ƞ_p_^2^ = 0.094, *p* = 0.032; **Supplementary Fig. 1D**).

For IL-10, GE2 tended to reduce levels compared to vehicle-treated subjects (main effect of treatment: F_(1,47)_ = 3.744, Ƞ_p_^2^ = 0.074, *p* = 0.059; **Supplementary Fig. 1E**). GE2 treatment significantly reduced IL-13 levels in the dorsal hippocampus compared to vehicle (main effect of treatment: F_(1,44)_ = 5.346, Ƞ_p_^2^ = 0.106, *p* = 0.025; **Supplementary Fig. 1F**). There was also a trend towards increased levels of IL-13 in WD compared to SD subjects (diet: F_(1,45)_ = 3.601, Ƞ_p_^2^ = 0.074, *p* = 0.064). GE2 treatment also led to a significant reduction in the levels of the chemokine CXCL1 compared to vehicle (main effect of treatment: F_(1,45)_ = 7.319, Ƞ_p_^2^ = 0.140, *p* = 0.010; **Supplementary Fig. 1G**).

TNFα levels were higher in GE2-treated WD-fed subjects compared to GE2-treated SD subjects (*p* = 0.079, Cohen’s *d* =1.067; treatment and sex interaction: F_(1,45)_ = 6.122, Ƞ_p_^2^ = 0.120, *p* = 0.017; **Supplementary Fig. 1H**). Lastly, we found that GE2 treatment reduced levels of IFNγ compared to vehicle in females (*p* = 0.075, Cohen’s *d* =0.855; treatment and sex interaction: F_(1,46)_ = 6.349, Ƞ_p_^2^ = 0.119, *p* = 0.015; **Supplementary Fig. 1I**), but not males (*p* = 0.213).

*Ventral hippocampus:* WD-fed rats receiving GE2 treatment had higher levels of IL-1β than SD fed rats receiving GE2, regardless of sex (treatment and diet interaction: *p* = 0.030, Cohen’s *d* =0.589; treatment and diet interaction: F_(1,45)_ = 5.459, Ƞ_p_^2^ = 0.108, *p* = 0.024; **Supplementary Fig. 2A**). There was a trend towards an increase in IL-6 after GE2 treatment in WD fed compared to SD fed in males (*p* = 0.059, Cohen’s *d* =1.509; sex, treatment and diet interaction: F_(1,47)_ = 4.978, Ƞ_p_^2^ = 0.096, *p* = 0.030; **Supplementary Fig. 2B**), but not females (*p* = 0.922). Similarly, we saw in increase in IFNγ in WD-fed GE2-treated subjects compared to SD fed receiving this treatment (*p* = 0.006, Cohen’s *d* =0.980; treatment and diet interaction: F_(1,46)_ = 4.213, Ƞ_p_^2^ = 0.084, *p* = 0.046; **Supplementary Fig. 2C**). In addition, we found that WD consumption in males led to higher levels of IFNγ compared to WD fed females (*p* = 0.024, Cohen’s *d* =0.605; sex and diet interaction: F_(1,46)_ = 7.748, Ƞ_p_^2^ = 0.144, *p* = 0.008) and SD fed males (*p* = 0.006, Cohen’s *d* =1.221). For TNFα we found that vehicle-treated females had lower levels of TNFα than vehicle-treated males (*p* = 0.018, Cohen’s *d* =1.213; sex and treatment interaction: F_(1,43)_ = 5.124, Ƞ_p_^2^ = 0.106, *p* = 0.029; **Supplementary Fig. 2D**). The main effect of sex was also significant (main effect of sex: F_(1,43)_ = 4.530, Ƞ_p_^2^ = 0.095, *p* = 0.039), as well as the main effect of diet (main effect of diet: F_(1,43)_ = 7.031, Ƞ_p_^2^ = 0.106, *p* = 0.011). Lastly, no significant main effects or interactions were found for CXCL1 in this area (all *p* ≥ 0.269).

*Amygdala:* In the amygdala we found that GE2-treated females on the SD had higher levels of IL-1β than all other groups (all *p* ≤ 0.047; sex, treatment and diet interaction: F_(1,39)_ = 7.461, Ƞ_p_^2^ = 0.161, *p* = 0.009; **Supplementary Fig. 3A**). This effect was also present for IL-4 levels and IL-10, in which elevated levels of IL-4 (all *p* ≤ 0.014; sex, treatment and diet interaction: F_(1,39)_ = 6.082, Ƞ_p_^2^ = 0.145, *p* = 0.019; **Supplementary Fig. 3B**) and IL-10 (all *p* ≤ 0.017 sex, treatment and diet interaction: F_(1,40)_ = 4.258, Ƞ_p_^2^ = 0.155, *p* = 0.046; **Supplementary Fig. 3C**) in SD-fed GE2-treated females was observed, compared to all other groups. Furthermore found that GE2 increased the levels of chemokine CXCL1 compared to vehicles in SD-treated subjects (*p* = 0.035, Cohen’s *d* =0.844; treatment and diet interaction: F_(1,39)_ = 3.580, Ƞ_p_^2^ = 0.125, *p* = 0.023; **Supplementary Fig. 3D**) and that GE2 treatment in SD-fed subjects resulted in higher levels of CXCL1 than in WD-fed subjects (*p* = 0.034, Cohen’s *d* =0.424).

*PFC:* In the PFC, GE2 treatment led to a trend towards reduction in IL-1β levels in GE2 treated subjects compared to vehicle (main effect of treatment: F_(1,48)_ = 3.528, Ƞ_p_^2^ = 0.068, *p* = 0.066; **Supplementary Fig. 4A**). For IL-4, we found no effects of treatment, we did however find that the levels of this cytokine were higher in females than males (main effect of sex: F_(1,48)_ = 6.954, Ƞ_p_^2^ = 0.127, *p* = 0.011; **Supplementary Fig. 4B**). A similar trend was also seen for IL-10, although the sex effect did not reach significance (main effect of sex: F_(1,47)_ = 3.847, Ƞ_p_^2^ = 0.076, *p* = 0.056; **Supplementary Fig. 4C**). We found no significant effects or differences between treatment, diet groups and sex for IL-13 (*p* ≥ 0.163; **Supplementary Fig. 4D**). We found higher levels of IFNγ in WD-fed subjects compared to SD fed (main effect of diet: F_(1,47)_ = 6.785, Ƞ_p_^2^ = 0.126, *p* = 0.012; **Supplementary Fig. 4E**). We also found a trend towards elevated levels of IFNγ in females compared to males (main effect of sex: F_(1,47)_ = 3.211, Ƞ_p_^2^ = 0.064, *p* = 0.080). We also found elevated levels of chemokine CXCL1 in females compared to males (main effect of sex: F_(1,45)_ = 5.495, Ƞ_p_^2^ = 0.109, *p* = 0.024; **Supplementary Fig. 4F**) . Furthermore, WD exposure led to increased levels of CXCL1 compared to SD (main effect of treatment: F_(1,45)_ = 5.288, Ƞ_p_^2^ = 0.105, *p* = 0.026; **Supplementary Fig. 4F**). No other main effects of significant interactions were present for CXCL1 (*p* ≥ 0.093). In contrary to CXCL1, we found that WD led to reduced levels of TNFα compared to SD (main effect of diet: F_(1,49)_ = 3.760, Ƞ_p_^2^ = 0.071, *p* = 0.058; **Supplementary Fig. 4G**). There were no other significant main effects of interactions (all *p* ≥ 0.136).

**Results – Including individual treatment with GLP-1 and E2**

*GLP-1 and GE2 reduce body weight in WD fed males and females*

In SD-fed subjects, only GE2 treatment was sufficient to reduce body weight (SD VEH vs SD GE2: *p* = 0.029, Cohen’s *d* = 1.158; Treatment x Diet interaction: F_(3, 96)_ = 4.904, Ƞ_p_^2^ = 0.133, *p* = 0.003; **Supplementary Fig. 6A**). However, in WD-fed rats, weight-loss was present after GLP-1 (*p* = 0.000, Cohen’s *d* = 2.528), E2 (*p* = 0.024, Cohen’s *d* = 1.438) and GE2 (*p* = 0.000, Cohen’s *d* = 3.727) treatment. Treatment with GE2 led to significantly more weight-loss than E2 alone (*p* = 0.003, Cohen’s *d* = 1.546).

GE2 treatment led to weight-loss in both males (*p* = 0.000, Cohen’s *d* = 3.452) and females (*p* = 0.026, Cohen’s *d* = 1.114; Sex x Treatment interaction: F_(3, 96)_ = 5.142, Ƞ_p_^2^ = 0.138, *p* = 0.002), although weight-loss was greater in males than females (*p* = 0.029, Cohen’s *d* = 0.899). In males, E2 (*p* = 0.001, Cohen’s *d* = 3.225) and GLP-1 (*p* = 0.000, Cohen’s *d* = 2.818) alone also led to significant weight-loss, and treatment with these hormones led to greater weight-loss in males than females (E2: *p* = 0.024, Cohen’s *d* = 1.194; GLP-1: *p* = 0.004, Cohen’s *d* = 0.891).

The main effects of sex (F_(1, 96)_ = 16.833, Ƞ_p_^2^ = 0.149, *p* = 0.000), treatment (F_(3, 96)_ = 25.835, Ƞ_p_^2^ = 0.447, *p* = 0.000) and diet (F_(1, 96)_ = 23.861, Ƞ_p_^2^ = 0.199, *p* = 0.000) were also significant.

*WD significantly increases IWAT and GWAT and GLP-1-E2 did not significantly reduce these adipose depots*

As expected, WD significantly increased IWAT compared to SD (main effect of diet: F_(1, 95)_ = 15.189, Ƞ_p_^2^ = 0.138, *p* = 0.000; **Supplementary Fig. 6B**). Males had significantly more IWAT than females (main effect of sex: F_(1, 95)_ = 28.676, Ƞ_p_^2^ = 0.232, *p* = 0.000). GLP-1 treatment resulted in reduced IWAT compared to control (*p* = 0.012, Cohens *d* = 0.795; main effect of treatment (F_(3, 95)_ = 5.196, Ƞ_p_^2^ = 0.141, *p* = 0.002). GLP-1 (*p* = 0.009, Cohens *d* = 0.783) and E2 (*p* = 0.031, Cohens *d* = 0.543) treated rats had lower levels of IWAT than GE2. There were no significant interactions between these factors (all *p*’s > 0.315).

WD also led to increased levels of GWAT compared to SD (main effect of diet: F_(1, 93)_ = 25.887, Ƞ_p_^2^ = 0.218, *p* = 0.000; **Supplementary Fig. 6C**). Males had higher percentage of GWAT than females overall (main effect of sex: F_(1, 93)_ = 8.108, Ƞ_p_^2^ = 0.080, *p* = 0.005). There was a trend towards a significant main effect of treatment, however this did not reach significance (*p* = 0.055) and there were no significant interactions between these factors (all *p*’s > 0.167).

*GE2 treatment did not significantly reduce blood glucose levels*

E2- (vehicle: *p* = 0.006; Cohen’s *d* = 1.014; GE2: *p* = 0.010; Cohen’s *d* = 0.787) and GLP-1- (vehicle: *p* = 0.001; Cohen’s *d* = 1.129; GE2: *p* = 0.003; Cohen’s *d* = 0.920) treated rats had significantly lower basal blood glucose compared to vehicle and GE2 (main effect of treatment: F_(3, 95)_ = 7.752, Ƞ_p_^2^ = 0.197, *p* = 0.000; **Supplementary Fig. 6D**) for basal blood glucose levels after fasting, in which. The main effects of sex and diet were not significant, nor was there a significant interaction between these factors (all *p*’s > 0.163). In the GTT we found no significant main effects or interactions (all *p*’s > 0.109; **Supplementary Fig. 6E**).

*GLP-1-E2 treatment increases plasma PYY levels regardless of diet*

WD-fed rats had lower levels of GLP-1 than SD (main effect of Diet: F_(1, 95)_ = 5.334, Ƞ_p_^2^ = 0.053, *p* = 0.023; **Supplementary Fig. 7A**). There were no other significant main effects or interactions (all *p* ≥ 0.097). Next, we looked at glucagon and insulin, hormones regulated by GLP-1. As for glucagon, there was a trend towards higher levels in females, although this did not reach significance (main effect of Sex: *p* = 0.089; **Supplementary Fig. 7B**). Plasma insulin levels, on the other hand, were higher in males (main effect of Sex: F_(1, 83)_ = 5.416, Ƞ_p_^2^ = 0.061, *p* = 0.022; **Supplementary Fig. 7C**). E2 and GLP-1 treatment reduced insulin levels compared to both vehicle and GE2 (VEH vs E2: *p* = 0.043; Cohen’s *d* = 0.896; VEH vs GLP-1: *p* = 0.032; Cohen’s *d* = 0.766; E2 vs GE2: *p* = 0.006; Cohen’s *d* = 1.042; GLP-1 vs GE2: *p* = 0.007; Cohen’s *d* = 0.932; main effect of Treatment: F_(3, 83)_ = 4.210, Ƞ_p_^2^ = 0.132, *p* = 0.008). Leptin levels were also higher in males (main effect of Sex: F_(1, 95)_ = 14.349, Ƞ_p_^2^ = 0.131, *p* < 0.001; **Supplementary Fig. 7D**), and WD-fed rats had higher levels than SD-fed rats (main effect of Diet: F_(1, 95)_ = 15.911, Ƞ_p_^2^ = 0.143, *p* < 0.001). There were no significant interactions (all *p* ≥ 0.225). Plasma PYY levels were increased after GE2-treated compared to vehicle-treated rats (*p* = 0.007; Cohen’s *d* = 1.068; main effect of Treatment: F_(3, 79)_ = 4.863, Ƞ_p_^2^ = 0.156, *p* = 0.004; **Supplementary Fig. 7E**). Levels of PYY were also increased in GE2-treated rats compared to those receiving GLP-1 or E2 alone (GLP-1 vs GE2: *p* = 0.006; Cohen’s *d* = 0.812; E2 vs GE2: *p* = 0.005; Cohen’s *d* = 0.901). There was a trend towards an interaction between Treatment, Diet and Sex, but this did not reach significance (*p* = 0.057), and there were no other significant main effects or interactions (all *p* ≥ 0.240). Plasma ghrelin and c-peptide were also measured but the majority of the samples were below detection level and could therefore not be analyzed.

*GLP-1 and GE2 treatment increase freezing in the fear conditioning task in both sexes*

In the contextual fear conditioning task, vehicle- and E2-treated subjects presented with less freezing than GLP-1 and GE2 (VEH vs GLP-1: *p* < 0.001; Cohen’s *d* =2.241; VEH vs GE2: *p* < 0.001; Cohen’s *d* =1.961; E2 vs GLP-1: *p* < 0.001; Cohen’s *d* =1.636; E2 vs GE2: *p* < 0.001; Cohen’s *d* =1.444; main effect of treatment: (F_(3, 95)_ = 17.944, Ƞ_p_^2^ = 0.362, *p* < 0.001; **Supplementary** **Fig. 8A**). These effects were also present in the cued fear conditioning task (VEH vs GLP-1: *p* = 0.009; Cohen’s *d* =0.988; VEH vs GE2: *p* = 0.001; Cohen’s *d* =1.324; E2 vs GLP-1: *p* = 0.006; Cohen’s *d* =1.222; E2 vs GE2: *p* < 0.001; Cohen’s *d* =1.628; main effect of treatment: (F_(3, 81)_ = 8.313, Ƞ_p_^2^ = 0.235, *p* < 0.001; **Supplementary** **Fig. 8B**).There were no other significant main effects or interactions (all *p* ≥ 0.237).

*GE2 treatment reduces the levels of several cytokines in the dorsal hippocampus*

Significant results for individual cytokines in each brain region measured are presented under each figure. In brief, GE2 treatment reduced the levels of several cytokines in the dorsal hippocampus (**Supplementary Fig. 9**), an effect which was not present in rats treated with E2 or GLP-1 alone. In the ventral hippocampus, many cytokines were below detection level, however, for IL-1β and TNFα GE2 treatment led to similar cytokine levels to control whereas rats treated with E2 or GLP-1 alone presented with higher levels of these cytokines (**Supplementary Fig. 10**). In the amygdala, GE2 treatment led to an increase in several cytokines compared to vehicle, especially in SD-fed females, whereas GLP-1 and E2 treatment alone maintained similar levels to vehicle (**Supplementary Fig. 11**). Lastly, in the PFC, GE2 treatment resulted in similar levels to vehicle whereas GLP-1 and E2 treatment alone appear to reduce the levels of several cytokines in this brain region (**Supplementary Fig. 12**).

*GLP-1-E2 modulates PSD95 levels in a brain region- and sex-specific manner*

In the dorsal hippocampus we found a slight increase in PSD95 after GE2 treatment compared to vehicle (*p* = 0.015, Cohen’s *d* =0.305; F_(3, 93)_ = 8.048, Ƞ_p_^2^ = 0.206, *p* < 0.001; **Supplementary** **Fig. 13A**). GE2 treatment also led to higher levels of PSD95 than E2 alone (*p* < 0.001, Cohen’s *d* =0.604) and GLP-1 alone (*p* < 0.001, Cohen’s *d* =0.033), which appears to be driven mainly by males. There were no significant interactions between factors (all *p*’s > 0.065).

As for the ventral hippocampus, a certain subset of the samples were compromised during processing resulting in a smaller number of samples included in the analysis. In the remaining samples we observed that GLP-1 and E2 led to lower levels of PSD95 than vehicle and GE2 (VEH vs E2: *p* = 0.004, Cohen’s *d* =1.393; VEH vs GLP-1: *p* = 0.007, Cohen’s *d* =1.159; E2 vs GE2: *p* < 0.001, Cohen’s *d* =1.224; GLP-1 vs GE2: *p* = 0.001, Cohen’s *d* =1.241; F_(3, 69)_ = 8.065, Ƞ_p_^2^ = 0.260, *p* < 0.001; **Supplementary** **Fig. 13B**).

In the amygdala GLP-1 treatment led to lower levels of PSD95 (*p* = 0.020, Cohen’s *d* =0.766) whereas GE2 increased PSD95 (*p* = 0.009, Cohen’s *d* =0.890; main effect of treatment: F_(3, 94)_ = 4.322, Ƞ_p_^2^ = 0.121, *p* = 0.007; **Supplementary** **Fig. 13C**). In the prefrontal cortex there were no significant main effects (all *p*’s > 0.666, **Supplementary Fig. 13D**), or interactions between the factors (all *p*’s > 0.402).
